# Hypocretin/orexin neurons encode social discrimination and exhibit a sex-dependent necessity for social interaction

**DOI:** 10.1101/2022.08.19.504565

**Authors:** Matthew Dawson, Dylan J Terstege, Naila Jamani, Mio Tsutsui, Dmitrii Pavlov, Raluca Bugescu, Jonathan R Epp, Gina M Leinninger, Derya Sargin

**Author notes:** **Corresponding author:** Derya Sargin Address: University of Calgary 2500 University Ave Dr NW T2N 1N4 Calgary, AB, Canada.

## Abstract

Intraspecies social interactions are integral for survival and maintenance of society among all mammalian species. Yet, our understanding of the neural systems and mechanisms involved in the establishment of social connectedness are limited. Since their initial discovery as regulators of sleep/wakefulness and appetite in the brain, the hypocretin/orexin neurons have also been shown to play an essential role in modulating energy homeostasis, motivated and emotional behavior. These neurons are located exclusively in the hypothalamus which, regulates complex and goal-directed behaviors. The hypothalamus also plays an important role in the modulation of social behavior by encoding internal states. However, our understanding of the role of hypocretin neurons in social behavior is currently limited. To address this knowledge gap, we performed a combination of fiber photometry and machine learning based behavioral analysis in female and male mice expressing GCaMP6s in hypocretin neurons. We then applied optogenetic and pharmacological inhibition of hypocretin neuron signaling to determine the necessity of the hcrt neuron population for social behavior. Our results indicate that hypocretin neurons exhibit a robust increase in activity in response to social interaction in both female and male mice. We show here for the first time a social discrimination signal that is encoded differentially by hcrt neurons based on the nature of the social encounter. The intensity of the hcrt neuron activity predicts the subsequent duration of social interaction. The optogenetic inhibition of hypocretin neuron activity during social behavior leads to a reduction in the amount of time mice are engaged in social interaction in males but not in females. Blocking hcrt1 (orexin 1) receptors similarly reduces social interaction in males only. Reduced hcrt1 receptor signaling results in increased activity in the insular cortex and reduced activity in the VTA after social interaction in male mice. Together, these data implicate the lateral hypothalamus hypocretin neurons as a sexually dimorphic key regulator within the larger network of neural systems involved in social behavior. Our findings carry significant implications for the treatment of neuropsychiatric diseases characterized by social dysfunction, particularly considering the varying prevalence observed across different sexes.

## Introduction

Social behaviors are an essential component of survival for most organisms. Effectively and appropriately communicating with members of one’s species enables an individual to gain access to information about foraging opportunities, potential mates and detection of threats. Interactions between conspecifics also provide the foundation for social group hierarchies, providing members with protection from predation, support from others within the group, and obtaining emotional rewards. The importance of these gains for the success of an organism has led to lasting evolutionary conservation of social behaviors, over time reinforcing the development of complex neural systems for the more sophisticated and refined methods of social interaction we see in mammals.

For organisms to interact socially, each must collect and process sensory signals from their counterparts and their environment, integrate this information with their internal state (e.g., memories of prior experiences, affective state, evaluation of risks and rewards), determine the appropriate behavioral response for the situation, then perform a sequence of corresponding motor patterns. These processes must function in concert simultaneously and rapidly to accommodate the highly dynamic and unpredictable process of social interaction (Chen and Hong, 2018). The resulting neural networks required to orchestrate this are vast, and we are still early in our understanding of which anatomical regions of the brain are involved and what their individual contribution to social behaviors is. Despite the challenge inherent to elucidating the underlying mechanisms of such complicated behaviors, many recent studies have made significant advances in our understanding of the neural substrates of social behaviors. Research in rodent models has demonstrated the involvement of numerous cortical and subcortical regions in production and direction of social behavior (Anderson, 2016; Clancy et al., 1984; Gunaydin et al., 2014; Hashikawa et al., 2016; Y. Hashikawa et al., 2017; Hofer, 1996; Hong et al., 2014; Keller et al., 2006; Kingsbury et al., 2019; Li et al., 2016; Marlin et al., 2015; Okuyama, 2018; Takahashi et al., 2014; Unger et al., 2015; Veening and Coolen, 2014; Wei et al., 2021). These regions have been found to form extensive functional circuits between each other and work to govern the array of parallel processes required during social interaction.

The various regions of the hypothalamus play a particularly important role in the modulation of social behavior by encoding internal states (Lo et al., 2019). The evolutionarily conserved hypothalamus (Xie and Dorsky, 2017) regulates critical survival functions by processing sensory and neuroendocrine signals and producing internal states to promote the organism to adjust its behavior to maintain homeostasis (Saper and Lowell, 2014). Certain internal states including emotion, memories of past experiences, motivation, and arousal are critical mediators in social behaviors as they determine the appropriate behavioral expression in response to incoming sensory information (Anderson, 2016). The modulatory power of these internal states over the social behavior of an animal is immense, to the degree that identical sensory stimuli and environmental conditions can elicit completely different behavioral responses based on the animal’s internal state (Chen and Hong, 2018). A shared underlying characteristic to all these internal states is that behavioral arousal is required to produce attention to socially relevant signals and motivation to investigate and interact with the conspecific from which they originate. This systemic arousal directed towards social stimuli is a process central to social behavior and closely overlaps with the functional role of the lateral hypothalamus (LH). Since their discovery (de Lecea et al., 1998; Sakurai et al., 1998), the LH hypocretin (hcrt)/ orexin neurons have been shown to govern arousal (Bourgin et al., 2000; Del Cid-Pellitero and Garzón, 2011; Hasegawa et al., 2017, 2014; Yang et al., 2019), attentional (Fadel and Burk, 2010; Fadel and Frederick-Duus, 2008), and motivational (España et al., 2011, 2010; Fadel and Deutch, 2002; Vittoz et al., 2008) states which all contribute to modulation of social behavior. Hcrt neurons extensively project to and modulate the activity of brain regions involved in social interaction (Bisetti et al., 2006; Lungwitz et al., 2012; Samson et al., 2002; Sterley et al., 2018), social reward (Fadel and Deutch, 2002; Gunaydin et al., 2014; Payet et al., 2021; Wang et al., 2005), and social memory (Yang et al., 2013). However, their direct role in social behavior has been less studied.

To reveal how hcrt neuron activity contribute to social behavior, we employed Cre-inducible viral targeting combined with fiber photometry and optogenetic tools in *Hcrt^IRES-Cre^* mice during conspecific social interactions. We identified that hcrt neurons exhibit a robust increase in activity in response to social interaction. We demonstrate that hcrt neuron activity is more pronounced during interaction between unfamiliar mice compared to that observed during familiar interactions. In contrast, exposing mice to familiar or novel objects does not lead to a difference in the activity of hcrt neurons suggesting that differential activity is driven primarily by social interactions. We further found that social interaction recruits a larger proportion of hcrt neurons in males, compared with females. The acute optogenetic inhibition of hcrt neuron activity and reducing hcrt signaling by blocking hcrt-1 receptors during social behavior disrupts social interaction in male mice without an effect in females. In males, we detected increased insular cortex and reduced VTA activity in mice injected with the hcrt-1 antagonist and subjected to social interaction. Together, our data compelling evidence for the sex-dependent role for hcrt neurons and downstream postsynaptic regions in social behavior.

## Results

### The activity of the LH hcrt neuron population increases in response to social investigation

To monitor the activity of hcrt neuron population in response to social approach and investigation, we used fiber photometry while performing a 3-chamber sociability test (Moy et al., 2004). We infused a Cre-dependent adeno-associated virus (AAV) encoding the fluorescent sensor GCaMP6s into the LH of *Hcrt^IRES-Cre^* mice. We found that >90 % of virus-labeled cells co-expressed Hcrt, while 58 % Hcrt+ cells were transduced with the virus (**Figure 1A, B; Figure S1, S2**), similar to the previously published *Hcrt-cre* knock-in lines (Giardino et al., 2018; Mitchell et al., 2020). An optical fiber was implanted above the LH to allow the delivery of excitation light and collection of GCaMP6s fluorescence from hcrt neurons specifically (**Figure 1A, B**). After 5 min baseline recording in the middle zone of the 3-chamber apparatus, mice were given access to freely explore the chamber containing a confined social target (age and sex-matched novel conspecific) in one chamber and an empty non-social zone in other chamber (**Figure 1C**). We first averaged the hcrt Ca^2+^ signals in response to the social interaction that occurred within 2 sec of the social zone entrance and compared this with the Ca^2+^ signals in mice during their interaction with the empty cup in the non-social zone. 2 sec duration after entrance was selected restrict our analysis to the bouts in which mice directly visited the sniffing zones after entering the chamber. In female mice, the hcrt neuron activity increased after entering the social and non-social zones (t>0 sec) and during interaction in the sniffing zones however, the activity between social and non-social zones remained non-significant (two-way ANOVA; effect of time: F(2, 18) = 15.96, *p* < 0.001, effect of zone: F(1, 18) = 0.07, *p* = 0.79) (**Figure 1D, E**). The area under the curve of the averaged hcrt Ca^2+^ responses were similar whether the female mice entered the social zone or the non-social zone (paired t-test; *p* = 0.96) (**Figure 1F**). In male mice, the hcrt Ca^2+^ response started to increase within 1 sec prior to and reached maximum levels right after entrance into the social zone (**Figure 1G**). The hcrt Ca^2+^ response in male mice was significantly larger during and after entrance into the social zone when compared with the non-social zone (two-way ANOVA; effect of time: F(2, 24) = 12.86, *p* = 0.0002, effect of zone: F(1,24) = 11.2, *p* = 0.0027) (**Figure 1H**). The area under the curve of the averaged hcrt Ca^2+^ responses was significantly larger when male mice entered the social zone rather than the non-social zone (*paired t-test; *p* = 0.025) (**Figure 1I**).

**Figure 1.**
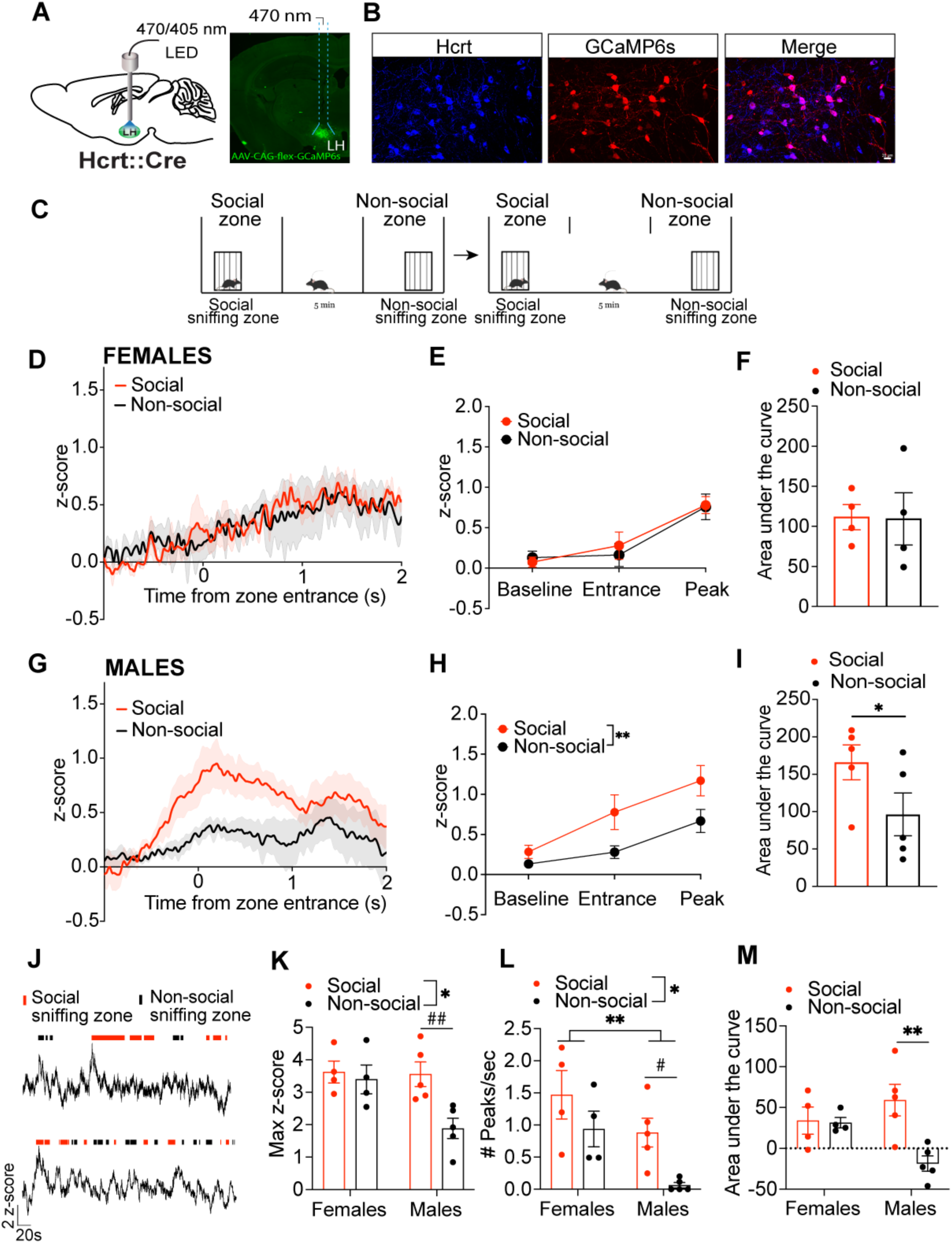
Increase in hcrt neuron activity in response to social investigation. **A)** Schematic (left) and a representative image (right) showing viral targeting and expression of GCaMP6s in LH hcrt neurons. **B)** Hypocretin-1 (blue) and GCaMP6s (red) co-localization in the LH of a *Hcrt^IRES-Cre^* mouse infused with the AAV-CAG-Flex-GCaMP6s. Scale bar, 20 um. **C)** 3-chamber social interaction paradigm during fiber photometry. Social and non-social zones are the compartments of the chamber that harbor the social target or the empty cup, respectively. Sniffing zones are defined as the area around the perimeter of the holder zones. **D)** Averaged hcrt Ca^2+^ traces over time aligned to the time of entrance (t = 0 sec) into the social zone in female mice (n = 4 mice). **E)** The hcrt Ca^2+^ signal for each mouse showing baseline (-1 to 0 sec), entrance (0 sec) and peak response after the entrance into the social zone (0-2 sec). In female mice, the hcrt Ca^2+^ signal increased upon entrance into the social and non-social zones and the activity in each zone was comparable (two-way ANOVA; effect of time: F(2, 18) = 15.96, *p* < 0.001, effect of zone: F(1, 18) = 0.07, *p* = 0.79). **F)** In female mice, the area under the curve of the averaged hcrt Ca^2+^ responses after entrance into social and non-social zones did not differ (paired t-test; *p* = 0.96). **G)** Averaged hcrt Ca^2+^ traces over time aligned to the time of entrance (t = 0 sec) into the social zone in male mice (n = 5 mice). **H)** In male mice, the hcrt Ca^2+^ signal increased upon entrance into the social and non-social zones and the activity in social zone was significantly greater than activity in the non-social zone (two-way ANOVA; effect of time: F(2, 24) = 12.86, *p* = 0.0002, effect of zone: F(1,24) = 11.2, \*\**p* = 0.0027). **I)** In male mice, the area under the curve of the averaged hcrt Ca^2+^ responses after entrance into the social zone was significantly greater compared to the non-social zone (paired t-test; \**p* = 0.025). **J)** A representative hcrt Ca^2+^ photometry trace aligned to the time spent in social (red) and non-social (black) sniffing zones from a female (above) and a male mouse (below). **K)** The maximum hcrt Ca^2+^ signal during interaction with the social target was larger compared with that during non-social target interaction regardless of sex (two-way ANOVA; effect of zone: F(1, 14) = 6.55, \**p* = 0.023, effect of sex: F(1, 14) = 4.56, *p* = 0.051). This difference was significant only in males (Fisher’s LSD; social vs non-social: females, *p* = 0.69, males, ^##^*p* = 0.004). **L)** The frequency of activity (peaks/sec) during interaction with the social target was significantly larger compared to the non-social interaction. In females, the frequency of hcrt activity was larger during social and non-social interactions, compared with the male mice (two-way ANOVA; effect of zone: F(1, 14) = 7.77, *p* = 0.014, effect of sex: F(1, 14) = 9.15, \*\**p* = 0.009). Planned comparisons revealed a significant difference for males but not females (Fisher’s LSD; social vs non-social: females, *p* = 0.16, males, ^#^*p* = 0.024). **M)** The area under the curve of the hcrt Ca^2+^ signal in each sniffing zone, normalized by the duration of time spent in each zone was significantly larger during interaction with the social target compared with the non-social target in male mice only (two-way ANOVA; zone X sex interaction: F(1, 14) = 6.74, *p* = 0.021; Tukey’s multiple comparisons test: Social vs non-social sniffing zone; Males \*\**p* = 0.006, females *p* = 0.99). Data represent mean ± SEM. The complete statistical output is shown in Supplementary Table S1.

We next compared the difference in the activity of hcrt neuron population while mice were actively sniffing the confined conspecific (social sniffing zone) or the empty cup (non-social sniffing zone) (**Figure 1J**). During interaction with the social target, hcrt Ca^2+^ response was significantly larger regardless of sex (two-way ANOVA; effect of zone: F(1, 14) = 6.55, *p* = 0.023, effect of sex: F(1, 14) = 4.56, *p* = 0.051) when compared to the time spent in the non-social sniffing zone (**Figure 1K**). Further analysis of the data using planned comparisons between social and non-social sniffing zones revealed that hcrt Ca^2+^ response was significantly larger in the social sniffing zone, compared with the non-social sniffing zone in males, but not in females (Fisher’s LSD; social vs non-social: females, *p* = 0.69, males, *p* = 0.004) (**Figure 1K**). The frequency of activity (peaks/sec) during interaction with the social target was significantly higher compared to the non-social interaction, with females having an overall greater frequency (two-way ANOVA; effect of zone: F(1, 14) = 7.77, *p* = 0.014, effect of sex: F(1, 14) = 9.15, *p* = 0.009). Planned comparisons of the frequency between social and non-social sniffing zones revealed a significant difference for males but not females (Fisher’s LSD; social vs non-social: females, *p* = 0.16, males, *p* = 0.024) (**Figure 1L**). The area under the curve of the hcrt Ca^2+^ signal changes was significantly larger when male mice interacted with the social target compared to the interaction in the non-social sniffing zone. This difference was not significant in females (two-way ANOVA; zone X sex interaction: F(1, 14) = 6.74, *p* = 0.021; Tukey’s multiple comparisons test: Social vs non-social sniffing zone; Males \*\**p* = 0.006, females *p* = 0.99) (**Figure 1M**).

Taken together, our data show that the response of the hcrt neuron population is larger when mice are actively sniffing the social conspecific when compared with the non-social exploration of an empty cup. This response is more prominent in males when they enter the social zone and interact with the social target, compared to females. However, regardless of sex, mice show an overall higher hcrt activity during the time they spent actively interacting with the social target in the sniffing zone.

### Differential activity in the LH hcrt neuron population in response to familiar and novel social interaction

While the 3-chamber sociability test allows exploration of social interaction within a controlled setting, it does not allow direct social interaction. To determine the changes in the activity of hcrt neuron population during reciprocal and active social interaction, we performed fiber photometry in the resident mice while they are socially interacting in their home cage. Accordingly, we performed 5 min baseline recordings after which the resident mouse is first introduced to a cage partner (familiar conspecific) and allowed to interact for 5 min. After removing the familiar conspecific from the cage followed by a 5 min additional baseline recording, the resident mouse was introduced to a novel age-, sex-and strain-matched conspecific (stranger) and allowed to interact for 5 min (**Figure 2A**). This order was selected due to the possibility that potentially aggressive encounters with the stranger mice would increase the stress levels and confound subsequent interaction with a familiar mouse. In addition, the behavior of the familiar mouse towards the experimental mouse could be influenced by the odor of a previous stranger. To rule out the possibility of habituation, we determined changes in the hcrt neuron activity during subsequent interactions. We did not detect a difference in the hcrt neuron activity in the resident mouse when a familiar or a stranger mouse was introduced twice in a row (**Figure S3**). In the home cage, social interaction was initiated exclusively by the resident mouse approaching the mouse introduced into the cage (Fam: 1.41 ± 0.23 sec, Stranger: 1.6 ± 0.55 sec after introduction). Social approach was defined as the time the resident mouse initiated movement toward the social target (Gunaydin et al., 2014). Averaging the hcrt Ca^2+^ signals over time in both female and male mice showed that the hcrt activity in resident mice started to increase prior to initial approach, reached maximum levels right after the initial approach and sniffing and returned to baseline minutes after the first interaction (**Figure 2B, C**). The analysis of the area under the curve of the averaged hcrt Ca^2+^ traces showed that the hcrt Ca^2+^ response was significantly larger in female and male resident mice during interaction with the unfamiliar conspecifics compared to the interaction with the familiar cage mates (two-way ANOVA; effect of conspecific: F(1, 26) = 8.96, *p* = 0.006, effect of sex: F(1, 26) = 1.78, *p* = 0.19). Planned comparisons between familiar and stranger interactions revealed a significant difference in males but not in females (Fisher’s LSD; familiar vs stranger: females, *p* = 0.1, males, *p* = 0.016) (**Figure 2D, E**). The maximum hcrt activity (max z-score ΔF/F) within 10 sec of initial approach was significantly greater during interaction between unfamiliar (resident and stranger) compared to that observed during familiar interactions in both female and male mice, with activity being larger during both types of social interaction in male mice compared with female mice (two-way ANOVA; effect of conspecific: F(1, 26) = 8.01, *p* = 0.009, effect of sex: F(1, 26) = 8.29, *p* = 0.008). Planned comparisons revealed a significant difference in females (Fisher’s LSD; familiar vs stranger: females, *p* = 0.03, males, *p* = 0.09) (**Figure 2F**). We did not detect a difference in variances between females and males (Levene’s test; z-score; F(3, 26) = 1.11, *p* = 0.36). The social novelty index did not differ between the sexes (mean ± SEM: Males, 0.21 ± 0.08; Females, 0.22 ± 0.1; unpaired t-test, t = 0.32, df = 13, *p* = 0.97).

**Figure 2.**
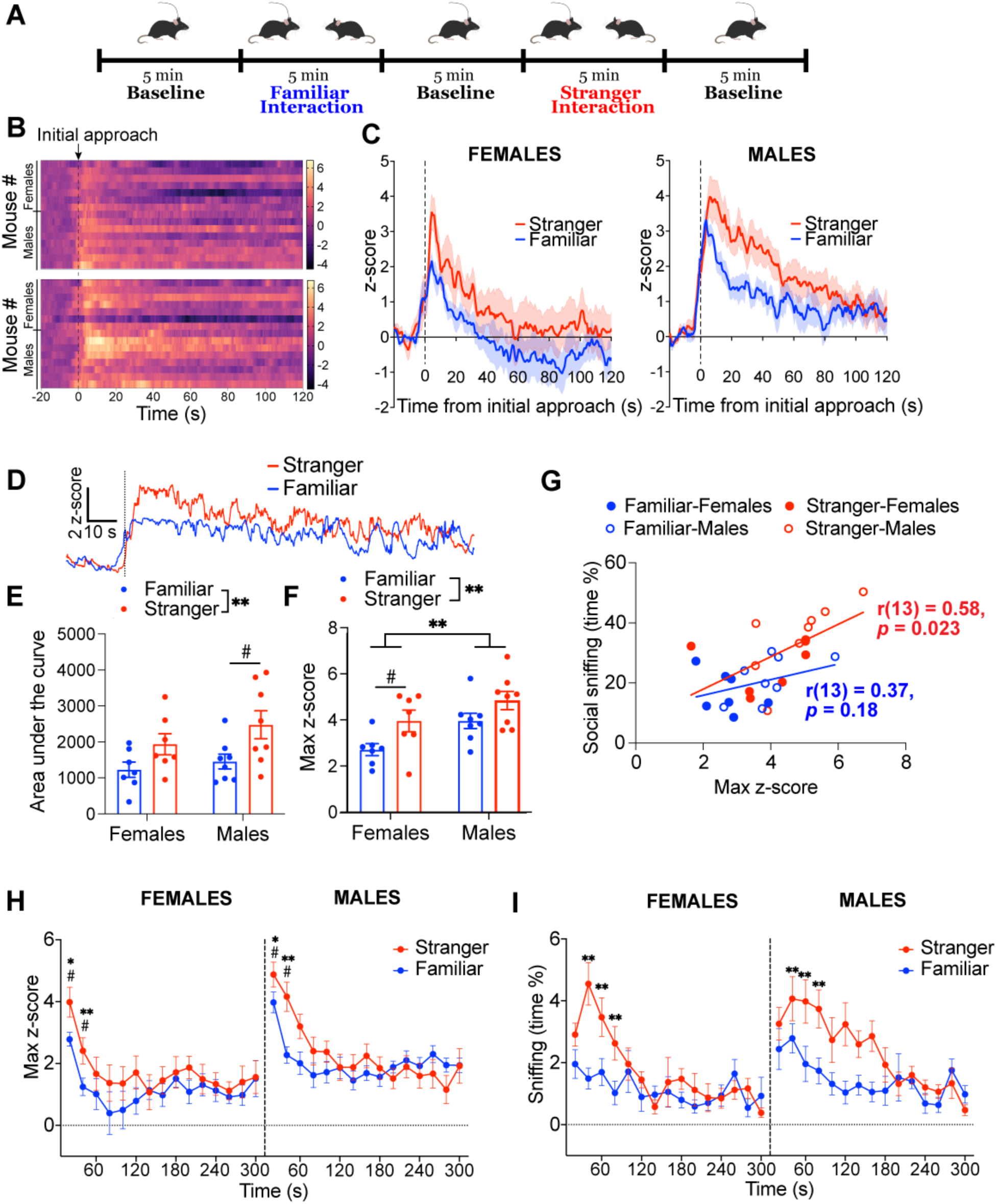
Differential hcrt neuron activity during social interaction in home cage. **A)** Experimental paradigm showing social interaction test during fiber photometry. After baseline recordings for 5 min in the home cage, the resident experimental mouse is introduced to a familiar (cage mate) followed by a stranger (novel) mouse. **B)** Heatmaps showing hcrt Ca^2+^ signal for each mouse 20 sec prior to and 2 min after being introduced to a familiar (top) or a stranger (bottom) mouse. Data are aligned to initial approach by the resident mouse. **C)** Averaged Ca^2+^ signal changes in hcrt neurons in female (n = 7) and male (n = 8) mice during interaction with a stranger or a familiar mouse. The hcrt neuron activity peaks right after initial approach initiated by the experimental mouse and returns to baseline within 2 min. **D)** Representative hcrt Ca^2+^ traces from an experimental mouse during interaction with a stranger or a familiar mouse. **E)** The area under the curve of the hcrt Ca^2+^ response within the first 2 min of social behavior is larger when the resident mice interact with a stranger compared with a familiar mouse (two-way ANOVA; effect of conspecific: F(1, 26) = 8.96, \*\**p* = 0.006, effect of sex: F(1, 26) = 1.78, *p* = 0.19; Fisher’s LSD; familiar vs stranger: females, *p* = 0.1, males, ^#^*p* = 0.016). **F)** The maximum hcrt Ca^2+^ signal in response to initial approach and sniffing is larger when the resident mice are introduced to a stranger compared to a familiar cage mate. The hcrt Ca^2+^ signal is larger in male mice compared to female mice during interaction with familiar and stranger conspecifics (two-way ANOVA; effect of conspecific: F(1, 26) = 8.01, \*\**p* = 0.009, effect of sex: F(1, 26) = 8.29, \*\**p* = 0.008; Fisher’s LSD; familiar vs stranger: females, ^#^*p* = 0.03, males, *p* = 0.09). **G)** Scatter plot showing the correlation between the maximum hcrt Ca^2+^ signal in the resident mice and the time they spent sniffing the stranger (r(13)=0.58, \**p*=0.023) or the familiar (r(13)=0.37, *p*=0.18) conspecific. **(H)** The maximum hcrt Ca^2+^ signal in the resident mice over time during social behavior is shown for both females and males. The greater hcrt activity right after first approach and sniffing decreased over time in both females and males (three-way ANOVA; time X conspecific interaction: F(14, 364) = 2.32, *p* = 0.004; Tukey’s multiple comparisons test: Stranger vs familiar at 20 sec \**p* = 0.035, 40 sec \*\**p* = 0.009, all other time points *p’s* > 0.05). Male mice showed a greater hcrt Ca^2+^ signal within the first minute of social behavior, compared with females (three-way ANOVA; time X sex interaction: F(14, 364) = 2.19, *p* = 0.007; Tukey’s multiple comparisons test: Male vs female at 20 sec ^#^*p* = 0.013, 40 sec ^#^*p* = 0.033, all other time points *p’s* > 0.05). **I)** Time spent sniffing by the resident mice over time during social behavior is shown for females and males. Mice spent more time sniffing the stranger compared to familiar, which declined over time (three-way ANOVA; time X conspecific interaction: F (14, 364) = 3.46, *p* < 0.001, Tukey’s multiple comparisons test: Stranger vs familiar at 40 sec \*\**p* < 0.001, 60 sec \*\**p* = 0.0013, 80 sec \*\**p* = 0.005 all other time points *p’s* > 0.05). Male mice engaged more in social interaction than females (three-way ANOVA; effect of sex: F(1, 26) = 4.62, *p* = 0.041). Data represent mean ± SEM. The complete statistical output is shown in Supplementary Table S2.

Next, we sought to determine the interaction between the initial large increase in hcrt activity with social behavior. We found a significant correlation between the first interaction-induced maximum hcrt activity and the time the resident mice spent sniffing the stranger mice (interaction %) within the subsequent 5 min of the social behavior task (Stranger: r(13) = 0.58, \**p* = 0.023, Familiar: r(13) = 0.37, *p* = 0.18) **(Figure 2G)**. To further determine the relation between the hcrt signal and social behavior, we plotted the maximum hcrt activity (**Figure 2H**) along with the duration of the time spent by the resident mice sniffing the conspecifics (**Figure 2I**) over time. These data have been analyzed by three-way ANOVA with sex and conspecific as factors and time as a repeated measure. The maximum hcrt activity was the greatest during the first minute after mice met for the first time and declined over the subsequent minutes of the task (effect of time: F(14, 364) = 21.26, *p* < 0.001). This was found to be dependent on both sex (time X sex interaction: F(14, 364) = 2.19, *p* = 0.007; Tukey’s multiple comparisons test: Male vs female at 20 sec *p* = 0.013, 40 sec *p* = 0.033, all other time points *p’s* > 0.05), and conspecific (time X conspecific interaction: F(14, 364) = 2.32, *p* = 0.004; Tukey’s multiple comparisons test: Stranger vs familiar at 20 sec *p* = 0.035, 40 sec *p* = 0.009, all other time points *p’s* > 0.05) (**Figure 2H, I**) suggesting the engagement of this neuron population during initial contact. The time spent sniffing was significantly greater in males than females (effect of sex: F(1, 26) = 4.62, *p* = 0.041). Both female and male mice spent a significantly greater time sniffing the stranger mice compared with familiar cage mates (effect of conspecific: F(1, 26) = 11.81, *p* = 0.002). Similar to the decrease in the hcrt activity, the initial difference in the time spent sniffing the stranger and familiar mice declined over time (time X conspecific interaction: F (14, 364) = 3.46, *p* < 0.001, Tukey’s multiple comparisons test: Stranger vs familiar at 40 sec *p* < 0.001, 60 sec *p* = 0.0013, 80 sec *p* = 0.005 all other time points *p’s* > 0.05).

The greater social interaction-induced hcrt activity increase in males may depend on a larger number of neurons infected, when compared with females. To determine this, we performed immunohistochemistry for GFP and quantified the number of hcrt cells expressing GCaMP6s in both sexes. We found no significant difference in the number of GFP+ hcrt neurons infused with AAV encoding GCaMP6s (unpaired t-test, t = 1.05, df = 10, *p* = 0.32) (**Figure S4A, B**). We next hypothesized that the amount of hcrt neurons recruited by the initial interaction may be different between females and males. To further investigate this, we perfused mice 90 min after social interaction with a stranger conspecific and analyzed the percentage of hcrt neurons expressing cFos in both sexes. We found that the amount of hcrt neurons expressing cFos after social interaction was significantly larger in male mice, compared with females (unpaired t-test, t = 3.56, df = 6, *p* = 0.012) (**Figure S4C, D**). In contrast, we did not detect a difference in the number of hcrt neurons between females and males (unpaired t-test, t = 0.34, df = 6, *p* = 0.74) (**Figure S4E**).

Overall, these data show that hcrt neuron activity plays a significant and differential role in social interaction. We provide the first evidence that unfamiliar mice evoke a greater response in hcrt neuron activity than familiar mice. The peak hcrt signal induced by the initial approach and sniffing determines the subsequent time of interaction between the unfamiliar animals. Both female and male mice show an increase hcrt activity in response to stranger interaction compared with familiar however, male mice show greater hcrt activity and sniffing behavior regardless of the conspecific. Moreover, males have a greater percentage of hcrt neurons that are activated in response to social interaction, as indicated by elevated cFos levels compared with females.

### Differential hcrt neuron activity during social interaction persists in the novel cage but absent in response to object investigation

The choice of the home cage social interaction assay for our experiments allowed us to assess social interaction within the territorial environment of the resident mice. To determine whether the greater hcrt signal to a stranger conspecific persists in a non-territorial novel setting, we placed the experimental mice in a cage with novel bedding and performed fiber photometry during interaction with familiar and stranger mice (**Figure 3A**). In a novel cage, mice spent significantly more time engaging in non-social behaviors including rearing, ambulation and grooming compared to those in the home cage (**Figure S5A**). However, the time mice spent interacting with the familiar or the stranger conspecific was comparable in both cages (**Figure S5B-S5D**).

**Figure 3.**
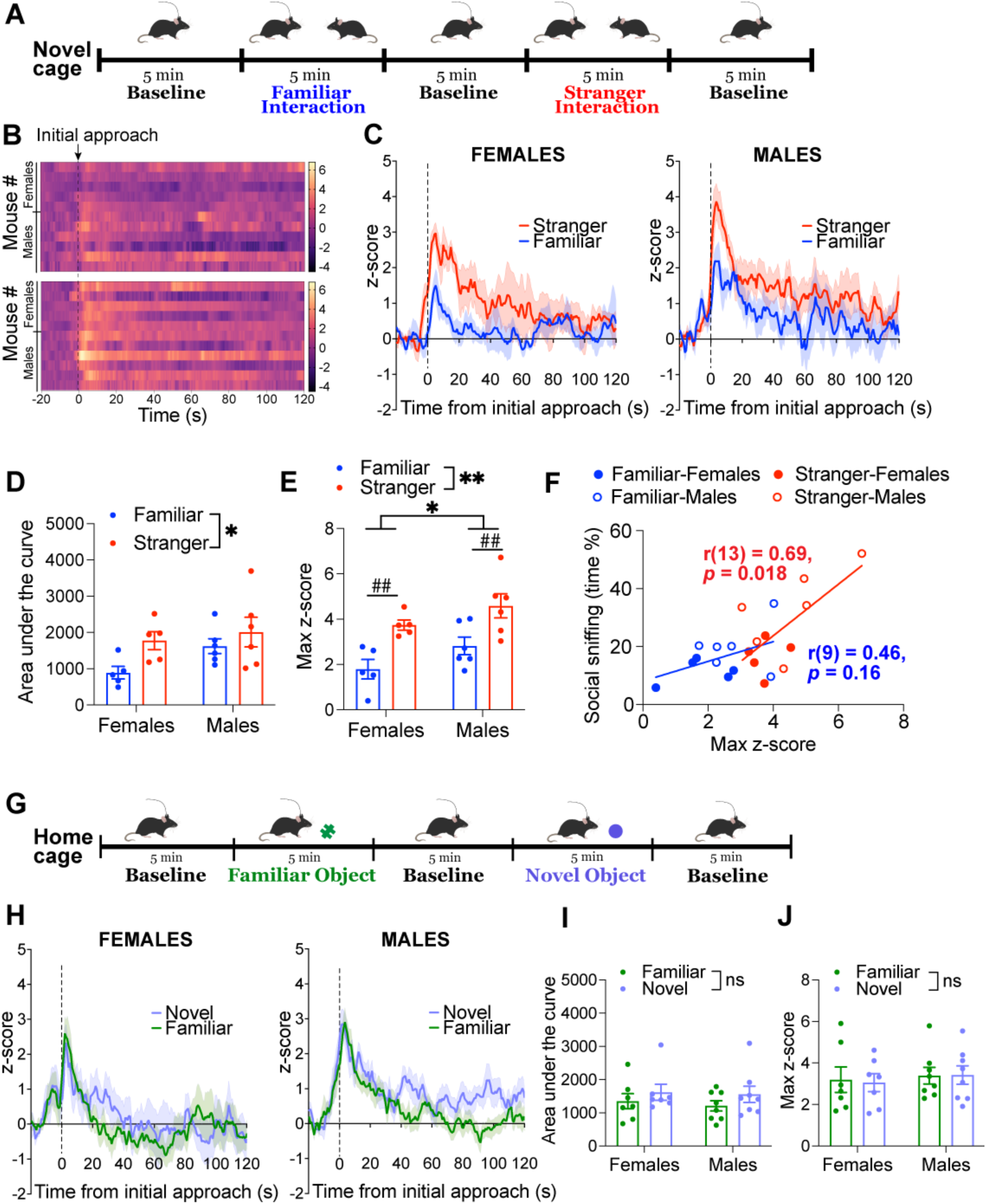
Differential hcrt activity during social interaction persists in a non-territorial setting and is absent during object investigation. **A)** Experimental paradigm showing social interaction in the novel cage test during fiber photometry. After baseline recordings for 5 min in a cage with novel bedding, the resident experimental mouse is introduced to a familiar (cage mate) followed by a stranger (novel) mouse. **B)** Heatmaps showing hcrt Ca^2+^ signal for each mouse 20 sec prior to and 2 min after being introduced to a familiar (top) or a stranger (bottom) mouse. Data are aligned to initial approach by the experimental mouse. **C)** Averaged Ca^2+^ signal changes in hcrt neurons in female (n = 5) and male (n = 6) mice during interaction with a stranger or a familiar mouse in the novel cage. The hcrt neuron activity peaks right after initial approach initiated by the experimental mouse and returns to baseline within 2 min. **D)** The area under the curve of the hcrt Ca^2+^ response within the first 2 min of social behavior is larger when the experimental mice interact with a stranger compared with a familiar mouse (two-way ANOVA; effect of conspecific: F(1, 18) = 4.91, \**p* = 0.038, effect of sex: F(1, 18) = 2.83, *p* = 0.11). **E)** The maximum hcrt Ca^2+^ signal in response to initial approach and sniffing is larger when the experimental mice are introduced to a stranger compared to a familiar cage mate. The hcrt Ca^2+^ signal is greater in male mice compared to female mice regardless of the social interaction type (two-way ANOVA; effect of conspecific: F(1, 18) = 19.14, \*\**p* = 0.0004, effect of sex: F(1, 18) = 4.87, \**p* = 0.041; Fisher’s LSD; familiar vs stranger: females, ^##^*p* = 0.006, males, ^##^*p* = 0.006). **F)** Scatter plot showing the correlation between the maximum hcrt Ca^2+^ signal in the experimental mice and the time they spent sniffing the stranger r(13)=0.69, \**p*=0.018) or the familiar (r(13)=0.46, *p*=0.16) conspecific. **G)** Experimental paradigm showing fiber photometry during object interaction test. After baseline recordings for 5 min in the home cage, the resident experimental mouse is introduced to a familiar or a novel object. **H)** Averaged Ca^2+^ signal changes in hcrt neurons in female (n = 7) and male (n = 8) mice during interaction with a familiar or a novel object in the home cage, aligned to the time of initial approach to the object. **I)** The area under the curve of the hcrt Ca^2+^ response within the first 2 min of object interaction is comparable when the experimental mice interact with novel or familiar objects (two-way ANOVA; effect of object: F(1, 26) = 1.97, *p* = 0.17, effect of sex: F(1, 26) = 0.21, *p* = 0.65). **J)** The maximum hcrt Ca^2+^ signal in response to initial approach to familiar and novel objects is comparable (two-way ANOVA; effect of object: F(1, 26) = 0.013, *p* = 0.91, effect of sex: F(1, 26) = 0.35, *p* = 0.56). \**p*<0.05, \*\**p*<0.01, ns; non-significant. Data represent mean ± SEM. The complete statistical output is shown in Supplementary Table S3.

Similar to our observations in the home cage, the increase in the hcrt Ca^2+^ signal reached maximum levels in female and male mice right after the first approach and sniffing in the novel cage (**Figure 3B, C**). The quantification of the area under the curve of the averaged hcrt Ca^2+^ traces revealed a greater hcrt activity when mice interacted with a stranger compared to a familiar mouse (two-way ANOVA; effect of conspecific: F(1, 18) = 4.91, *p* = 0.038) (**Figure 3D**). The maximum hcrt activity (max z-score ΔF/F) within 10 sec of initial approach was also significantly greater when mice interacted with a stranger compared with a familiar mouse (two-way ANOVA; effect of conspecific: F(1, 18) = 19.14, *p* = 0.0004). The hcrt activity was significantly larger in male mice during interaction with both familiar and stranger mice, when compared with females (two-way ANOVA; effect of sex: F(1, 18) = 4.87, *p* = 0.041). Planned comparisons between familiar and stranger interactions revealed a significant difference for both females and males (Fisher’s LSD; familiar vs stranger: females, *p* = 0.006, males, *p* = 0.006) (**Figure 3E**), and was comparable to the signal observed during interaction in the home cage (**Figure S5E, F**). Similar to the home cage setting, the initial interaction-induced maximum hcrt activity correlated with the amount of interaction with the stranger conspecifics (**Figure 3F**). We did not detect a difference in variances between females and males (Levene’s test; z-score; F(3, 18) = 3.32, *p* = 0.3). Social novelty index also did not differ females and males (mean ± SEM: Males, 0.17 ± 0.05; Females, 0.23 ± 0.07; unpaired t-test, t = 0.61, df = 9, *p* = 0.55). Altogether, these data show that the activity of hcrt neuron population is greater while mice are engaged with stranger conspecifics whether they are in their territorial home cage environment or in a novel cage, with higher hcrt activity and sniffing behavior in males in both settings.

To identify whether the differential activity in the hcrt neuron population is a general response to novelty or specific to social interaction, we performed fiber photometry while mice were exposed to a familiar or a novel object in their home cage. Experimental mice were pre-exposed to the familiar object for at least 3 days in their home cage (Okuyama et al., 2016). In this task, we presented the experimental mice with a familiar or a novel object following a 5 min baseline recording (**Figure 3G**). In both female and male mice, we observed an increase in the hcrt Ca^2+^ signal in response to initial approach and interaction with the objects (**Figure 3H**). In contrast to the differential hcrt signal observed during familiar and stranger mouse interactions, both the maximum hcrt Ca^2+^ signal and the area under the curve for the first 2 min of interaction were similar during novel and familiar object interactions (**Figure 3I, J**). This showed that the significant difference in the hcrt Ca^2+^ signal during social interaction was absent when mice interacted with the familiar or novel objects, suggesting that the differential activity is specific to social interaction.

Because female and male mice employ different strategies (Anderson, 2016; van den Berg et al., 2015), we further analyzed the two distinct modalities of social investigative behaviors that the experimental mice engaged in, including anogenital sniffing (examination of the anogenital area of the conspecific) and the head/torso sniffing (examination of the head and torso region of the conspecific (Lee et al., 2019; Sterley et al., 2018). Both female and male mice spent more time sniffing the anogenital and head/torso region of the stranger mice when compared with the familiar conspecifics, with males spending a greater time engaged in anogenital sniffing compared to females (Anogenital sniffing: two-way ANOVA; effect of conspecific: F(1, 26) = 18, \*\**p* = 0.002, effect of sex: F(1, 26) = 8.19, \*\**p* = 0.0082; Head/torso sniffing: two-way ANOVA; effect of conspecific: F(1, 26) = 3.17, *p* = 0.087, effect of sex: F(1, 26) = 0.99, *p* = 0.33) (**Figure S6A, B**). The analysis of the hcrt activity peaks time locked to each behavior yielded a significantly larger signal in female and male mice while they were engaged in the anogenital and head/torso sniffing of the stranger mice, compared to sniffing the familiar conspecifics (Anogenital sniffing: two-way ANOVA; effect of conspecific: F(1, 26) = 6.16, \**p* = 0.02, effect of sex: F(1, 26) = 3.62, *p* = 0.068; Head/torso sniffing: two-way ANOVA; effect of conspecific: F(1, 26) = 7.21, \**p* = 0.012, effect of sex: F(1, 26) = 2.6, *p* = 0.12) (**Figure S6C-S6F**). These data showed a stronger anogenital sniffing drive and a borderline increase in the activity of the hcrt neuron population during anogenital sniffing in male mice compared to females.

Taken together, our data showed that the activity of the LH hcrt neuron population increases in response to social or object investigation. However, while the larger hcrt neuron activity in response to novel social interaction indicates social discrimination, this difference in activity is absent when mice interact with familiar or novel objects. The sex differences with males responding to social interaction with larger anogenital sniffing and greater hcrt Ca^2+^ response compared with females are also absent when mice are interacting with objects.

### Inhibition of hcrt neuron activity disrupts social behavior in male mice

Having established that the activity of hcrt neuron population is highly responsive to social interaction, we next investigated whether hcrt neurons are required for social behavior. We infused a Cre-dependent adeno-associated virus (AAV) encoding EYFP (controls) or the blue light-activated chloride channel (iC++) (Berndt et al., 2016) into the LH of *Hcrt^IRES-Cre^*mice (**Figure 4A, B**). We validated that hcrt neurons expressing iC++ are inhibited in response to blue light stimulation using *ex vivo* electrophysiological recordings (**Figure S7A, B**). Because the first social interaction yields the maximum signal, which correlates with the subsequent interaction amount, we employed a photoinhibition protocol by silencing hcrt neurons in resident mice prior to the introduction of familiar or stranger conspecifics into the home cage (**Figure 4C**). The social interaction test was repeated on two separate days and the order of the conspecific was counterbalanced across different days. Social interaction was quantified as the summation of anogenital and head/torso sniffing. In female mice, the optogenetic inhibition of hcrt activity did not affect the time spent interacting with familiar or stranger conspecifics (two-way ANOVA; effect of group: F(1, 28) = 1.64, *p* = 0.21, effect of conspecific: F(1, 28) = 3.41, *p* = 0.07) (**Figure 4D**). In contrast, male mice spent significantly less time interacting with conspecifics when hcrt activity was inhibited (two-way ANOVA; effect of group: F(1, 24) = 13.82, *p* = 0.001, effect of conspecific: F(1, 24) = 2.66, *p* = 0.12; Fisher’s LSD; EYFP vs iC++: familiar, *p* = 0.07, stranger, *p* = 0.002) (**Figure 4E**). To verify that the optogenetic activation of iC++ induces inhibition of hcrt neurons effectively, we perfused a group of EYFP-and iC++-expressing mice 90 min after social interaction with a stranger conspecific. cFos immunostaining revealed a significant reduction in cFos+ hcrt neurons in female and male mice (two-way ANOVA; effect of group: F(1, 11) = 12.62, *p* = 0.004) (**Figure S7C, D**). We further found that male mice showed an overall greater hcrt cFos expression compared to females (two-way ANOVA; effect of sex: F(1, 11) = 6.78, *p* = 0.02) (**Figure S7C, D**).

**Figure 4.**
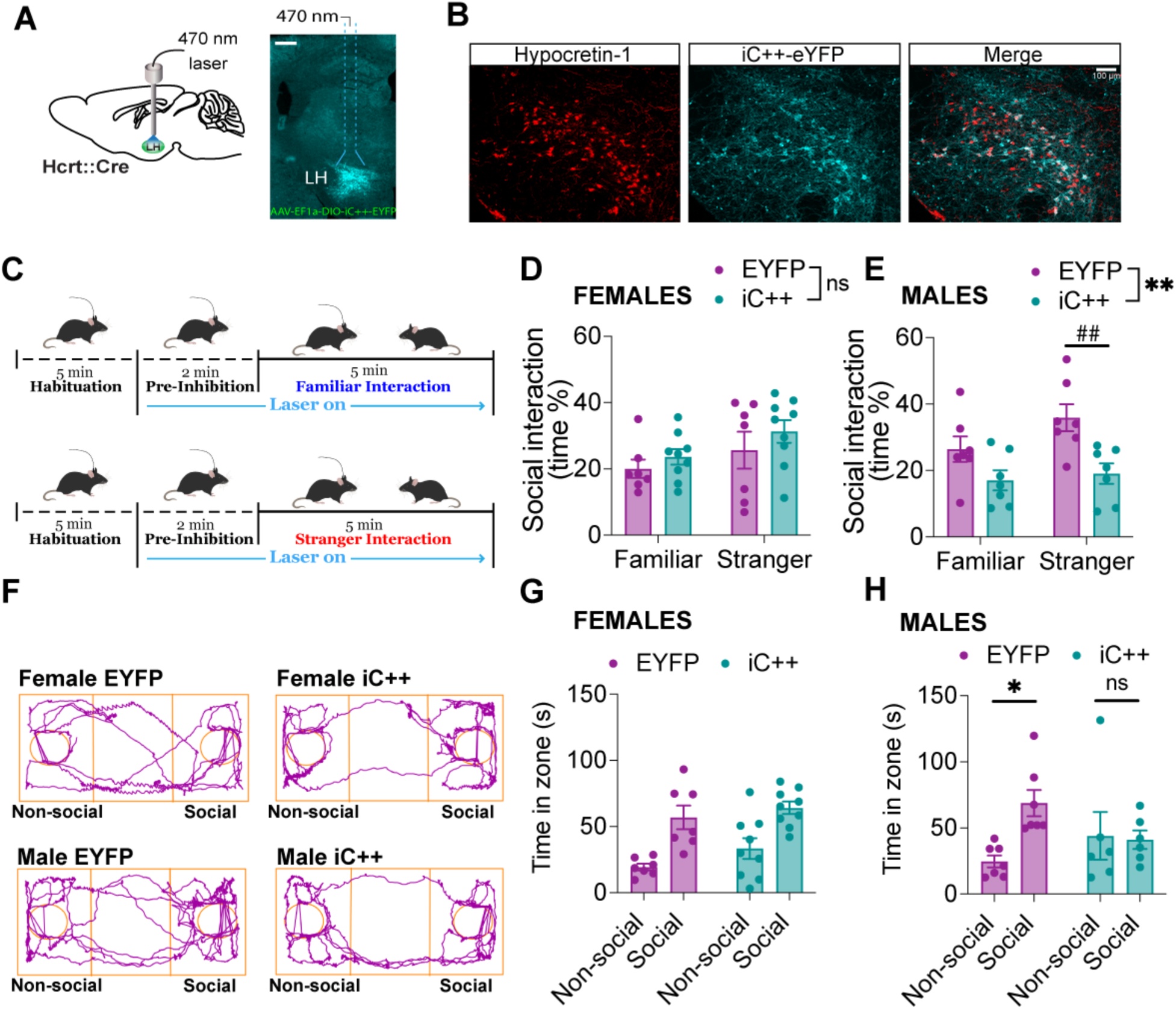
Acute optogenetic inhibition of hcrt neuron activity disrupts social interaction in male mice. **A)** Schematic (left) and a representative image showing viral targeting and expression of iC++ in LH hcrt neurons. **B)** Hypocretin-1 (red) and iC++ (cyan) co-localization in the LH of a *Hcrt^IRES-Cre^* mouse infused with the AAV-EF1a-DIO-iC++-EYFP. Scale bar, 100 um. **C)** Experimental paradigm showing 5 min social interaction with familiar or stranger mice after 5 min of fiber habituation and 2 min of hcrt neuron inhibition. Blue light was kept on during social interaction. Each interaction was performed on different days and the order of conspecific was counterbalanced. **D)** In female mice, optogenetic inhibition of hcrt neuron activity did not affect the time spent interacting with familiar or stranger conspecifics (two-way ANOVA; effect of group: F(1, 28) = 1.64, *p* = 0.21, effect of conspecific: F(1, 28) = 3.41, *p* = 0.07). **E)** Inhibition of hcrt neuron activity in male mice reduced interaction time with conspecifics, compared with EYFP mice (two-way ANOVA; effect of group: F(1, 24) = 13.82, \*\**p* = 0.001, effect of conspecific: F(1, 24) = 2.66, *p* = 0.12; Fisher’s LSD; EYFP vs iC++: familiar, *p* = 0.07, stranger, ^##^*p* = 0.002). **F)** Representative tracks from female and male EYFP and iC++ mice during in 3-chamber sociability task in the presence of blue light. **G)** Inhibition of hcrt neuron activity did not lead to a change in sociability in females in 3-chamber test (two-way ANOVA; effect of group: F(1, 28) = 2.59, *p* = 0.12, effect of zone: F(1, 28) = 26.56, *p* < 0.001). **H)** Male iC++ mice spent comparable time in the social and non-social sniffing zones of the 3-chamber in contrast to male EYFP mice (two-way ANOVA; group X zone interaction: F(1, 22) = 4.81, *\*p* = 0.039; Tukey’s multiple comparisons test: EYFP social vs non-social *p* = 0.029, iC++ social vs non-social *p* = 0.99). Data represent mean ± SEM. The complete statistical output is shown in Supplementary Table S4.

To determine whether sociability is affected similarly in a more controlled setting in which the stranger mouse is confined, we tested mice in a 3-chamber test while inhibiting hcrt neuron activity. While we detected no difference for sociability in females (two-way ANOVA; effect of group: F(1, 28) = 2.59, *p* = 0.12, effect of zone: F(1, 28) = 26.56, *p* < 0.001) (**Figure 4F, G**) and control males (**Figure 4F, H**), all showing a greater preference for the confined stranger, male iC++ mice spent comparable time in the social and non-social sniffing zones (two-way ANOVA; group X zone interaction: F(1, 22) = 4.81, *p* = 0.039; Tukey’s multiple comparisons test: EYFP social vs non-social *p* = 0.029, iC++ social vs non-social *p* = 0.99) (**Figure 4H**).

Because hcrt neuron activity is linked to locomotion (Karnani et al., 2020) and can be regulated by stress (Grafe et al., 2018, 2017; Sargin, 2019), we next investigated whether inhibition of hcrt neurons affects locomotor or anxiety-like behavior. We performed open field and elevated plus maze tests in the presence of blue light in EYFP-and iC++ expressing female and male mice. Acute inhibition of hcrt neuron activity during open field test did not lead to changes in distance traveled (two-way ANOVA; effect of group: F(1, 25) = 1.46, *p* = 0.24), or the time spent in the outer or the inner zone of the open field apparatus (two-way ANOVA; effect of group for time in outer zone: F(1, 25) = 1.45, *p* = 0.24, effect of group for time in inner zone: F(1, 25) = 1.45, *p* = 0.24) in iC++ female and male mice, compared with mice expressing EYFP (**Figure S8A-D**). In the open field test, both groups of female mice traveled a greater distance than male mice, indicating greater locomotor activity in females regardless of group (two-way ANOVA; effect of sex: F(1, 25) = 4.62, \**p* = 0.042). Female mice also spent more time in the outer zone (two-way ANOVA; effect of sex: F(1, 25) = 4.64, \**p* = 0.041) and less time in the inner zone of the open field (two-way ANOVA; effect of sex: F(1, 25) = 4.64, \**p* = 0.041), compared with male mice (**Figure S8A-D**). In the elevated plus maze, we did not detect a difference in locomotion (two-way ANOVA; effect of group: F(1, 23) = 0.64, *p* = 0.43, effect of sex: F(1, 23) = 3.02, *p* = 0.09) or anxiety-like behavior between the groups (two-way ANOVA; effect of group for time in closed arms: F(1, 23) = 1.21, *p* = 0.28, effect of group for time in open arms: F(1, 23) = 1.19, *p* = 0.28) or sexes (two-way ANOVA; effect of sex for time in closed arms: F(1, 23) = 1.07, *p* = 0.31, effect of sex for time in open arms: F(1, 23) = 0.67, *p* = 0.42) (**Figure S8E-H**).

Overall, acute optogenetic inhibition of hcrt neuron activity significantly reduced the time resident male mice spent interacting with the conspecifics, and decreased the preference of male mice for the social conspecific without affecting locomotor activity or anxiety-like behavior. In females, acute inhibition of hcrt neuron activity did not affect social interaction. These data suggest that hcrt neurons are necessary for normative social behavior in male mice while in females, the potential recruitment of different neural circuits may overcome the effects of reduced hcrt activity on social interaction.

### Reduced hcrt-1 receptor signaling alters the activity of the insular cortex and VTA and disrupts social interaction in male mice

Hcrt neurons are a group of heterogeneous cells that project to and modulate many different brain regions where they release hcrt and co-transmitters including dynorphin (Chou et al., 2001; Li and van den Pol, 2006) or glutamate (Schöne et al., 2012). To determine whether disrupted social interaction is dependent on hypocretin, we injected male and female mice with the hcrt-1 receptor antagonist SB-334867 or vehicle 30 min before social behavior (**Figure 5A**). Similar to the results of the optogenetic experiments, reduced hcrt-1 receptor signaling with SB-334867 administration significantly decreased social interaction in male mice only. SB-334867 did not lead to changes in the duration of time female mice engaged with the stranger conspecifics compared with the vehicle treatment however, females interacted with the stranger mice significantly less compared with male mice (two-way ANOVA; drug X sex interaction: F (1, 20) = 30.29, *\*\*p* < 0.001, Tukey’s multiple comparisons test: female vs male, vehicle *\*\*p* < 0.001, female vs male SB-334867 *p* = 0.53; female vehicle vs female SB-334867, *p* = 0.36, male vehicle vs male SB-334867, *\*\*p* < 0.001) (**Figure 5B**). To investigate the potential effects of SB-334867 on the locomotor activity or anxiety-like behavior, we performed open field right after vehicle and SB-334867 administrations and social behavior experiments (**Figure 5A**). The total distance traveled in the open field were not different between vehicle-and SB-334867-treated mice, with females traveling an overall greater distance compared to males (two-way ANOVA, distance traveled; effect of drug: F(1, 20) = 2.19, *p* = 0.15, effect of sex: F(1, 20) = 9.4, \*\**p* = 0.006) (**Figure 5C**). Moreover, SB-334867 administration did not yield any significant differences in the amount of time spent in the outer and inner zones of the open field (two-way ANOVA, time in outer zone; effect of drug: F(1, 20) = 2.1, *p* = 0.16, effect of sex: F(1, 20) = 1.7, *p* = 0.21, time in inner zone; effect of drug: F(1, 20) = 2.1, *p* = 0.16, effect of sex: F(1, 20) = 1.69, *p* = 0.21) (**Figure 5D, E)**. This suggested that changes in locomotor activity or anxiety levels did not contribute to the observed differences in social behavior.

**Figure 5.**
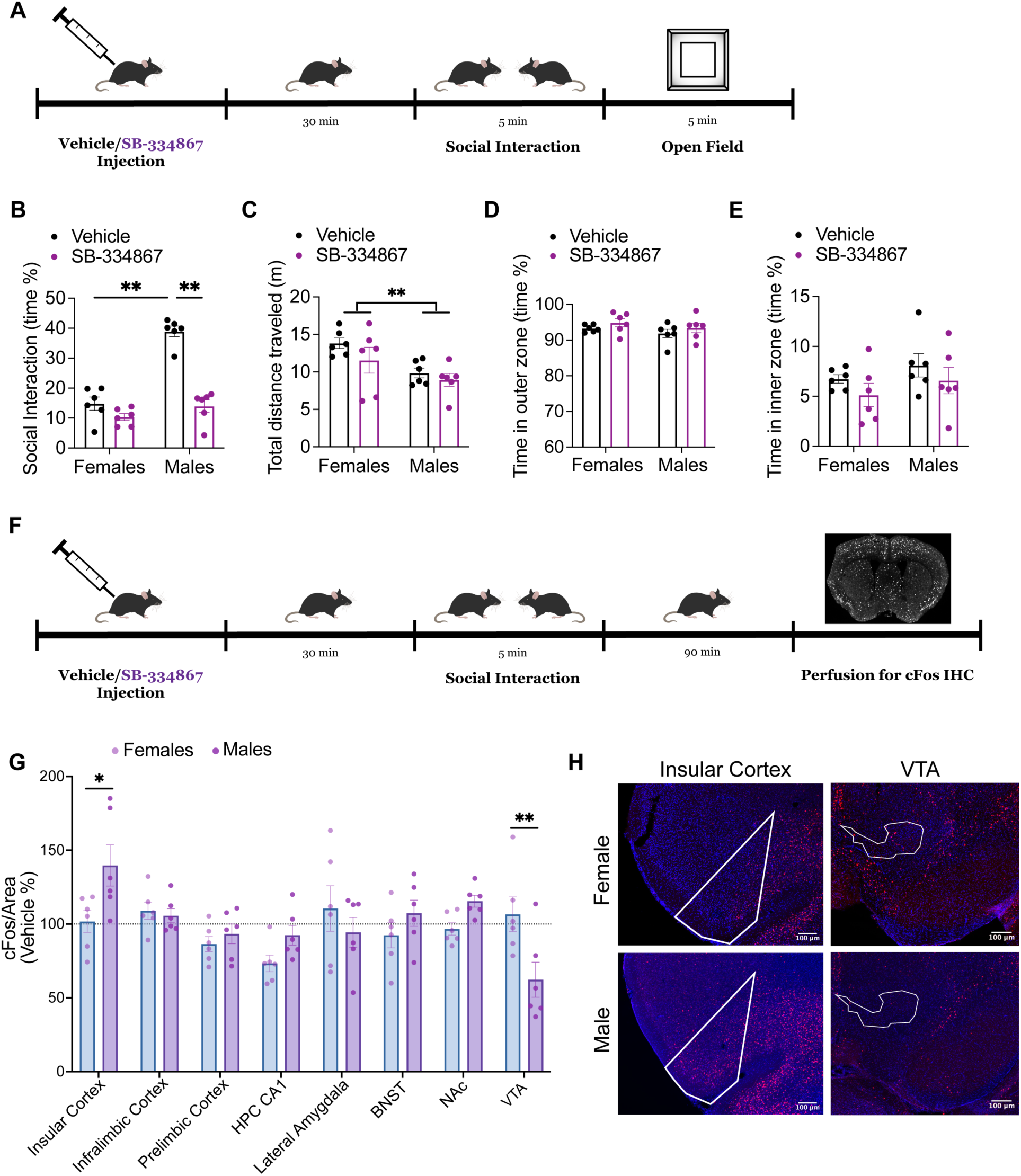
Blocking hcrt-1 receptors with SB334867 disrupts social interaction in male mice and is accompanied by alterations in the activity of the insular cortex and VTA. **A)** Experimental paradigm. Mice were first injected with either 30mg/kg of SB-334867 or vehicle. 30 min after injection, mice interacted with a novel conspecific for 5 min, followed by 5 min exploration in the open field. **B)** Injection of SB-334867 reduced social interaction with stranger conspecifics in male mice, without an effect in females (two-way ANOVA; drug X sex interaction: F (1, 20) = 30.29, *\*\*p* < 0.001, Tukey’s multiple comparisons test: Females vs males, vehicle *\*\*p* < 0.001, SB-334867 *p* = 0.53; Females vehicle vs SB-334867, *p* = 0.36, males vehicle vs SB-334867, *\*\*p* < 0.001). **C)** Total distance traveled is comparable between vehicle-and SB-334867-injected mice. Females traveled significantly greater distance in the open field, compared with males (two-way ANOVA; effect of drug: F(1, 20) = 2.19, *p* = 0.15, effect of sex: F(1, 20) = 9.4, \*\**p* = 0.006). The time spent in the outer zone **(D)** and the inner zone **(E)** of the open field is comparable between mice. **F)** Experimental paradigm. 30 min after vehicle or SB-334867 injection, mice were allowed to interact with a stranger conspecific for 5 min. Mice were perfused 90 min after social interaction. Brains were processed for cFos immunohistochemistry. **G)** Male mice injected with SB-334867 had significantly greater cFos density in the insular cortex and reduced cFos density in the VTA, compared with females injected with SB-334867 (two-way ANOVA; brain region X sex interaction: F(7, 80) = 4.12, *p* = 0.0007, Bonferroni’s multiple comparison test; males vs females, insular cortex \**p* = 0.02, VTA \*\**p* = 0.005). **H**) Representative images showing DAPI (blue) and cFos (red) immunostaining in the insular cortex and VTA of female and male mice.

We next investigated how social interaction in the presence or absence of hcrt-1 receptor blockade affects the activity of cortical and subcortical regions that are modulated by hcrt inputs and are implicated in social behavior. We perfused vehicle-and SB-334867-administered female and male mice 90 min after social interaction with a stranger conspecific and then immunolabeled the brains for cFos. We used the Flexible Atlas Segmentation Tool for Multi-Area Processing (FASTMAP) machine-learning pipeline (Terstege et al., 2022) for high-throughput quantification of cFos density in the insular, prelimbic, infralimbic cortical regions as well as the CA1 region of the hippocampus, lateral amygdala (LA), the bed nucleus of the stria terminalis (BNST), nucleus accumbens (NAc) and the ventral tegmental area (VTA). The comparison of the differences in cFos density between SB-334867-admnistered female and male mice revealed significant differences in two of the brain regions investigated. The density of cFos immunoreactive cells was significantly increased by SB-334867 treatment relative to control mice in the insular cortex and significantly decreased in the VTA of male mice, compared with females (two-way ANOVA; brain region X sex interaction: F(7, 80) = 4.12, *p* = 0.0007, Bonferroni’s multiple comparison; males vs females, insular cortex \**p* = 0.02, VTA \*\**p* = 0.005) (**Figure 5G, H**). Similarly, compared to the vehicle-injected mice, SB-334867-administered males showed significantly increased activity in the insular cortex and decreased activity in the VTA (two-way ANOVA; brain region X drug interaction: F(7, 80) = 3.18, *p* = 0.005, Bonferroni’s multiple comparison test; vehicle vs SB-334867, insular cortex \**p* = 0.016, VTA \*\**p* = 0.026) (**Figure S9A**). These changes were not observed in females (two-way ANOVA; effect of drug: F(1, 80) = 0.39, *p* = 0.53, effect of brain region: F(7, 80) = 0.94, *p* = 0.48) (**Figure S9B**). These data provide insights into the potential circuits that hcrt modulates differentially in females and males during social interaction.

## Discussion

Discrete neural subpopulations in the hypothalamus play a critical role in producing and modulating a variety of social behaviors, such as mating, aggression, and social bonding (Hashikawa et al., 2016; K. Hashikawa et al., 2017; Y. Hashikawa et al., 2017; Hull and Dominguez, 2007; Scott et al., 2015; Wu et al., 2014; Yoshihara et al., 2018). Similarly, research into the functional roles of hcrt neurons has begun to suggest that these cells are involved in the neural substrates of social processes, but the nature and extent of their involvement has not yet been determined. Here, using a combination of fiber photometry, optogenetic and pharmacological inhibition and social interaction assays, we characterized the role of hcrt neuron activity in social behavior. We demonstrate for the first time that the activity of hcrt neuron population is regulated by social approach and interaction and is critical for social behavior in a sex-dependent manner.

Photometry recordings revealed that hcrt neurons in female and male mice exhibit a robust increase in activity in response to social interaction. The elevation in activity was prominent during indirect social contact in the 3-chamber social interaction test or during reciprocal social interaction in the familiar home cage or less familiar novel cage. Moreover, hcrt neuron activity was more pronounced during interaction between unfamiliar mice compared to the increased activity observed during familiar interactions. These experiments provide the first evidence that the discrimination between familiar and unfamiliar conspecifics is differentially encoded by the hcrt neuron activity. In contrast, the increase in hcrt neuron activity remained the same when mice interacted with familiar or novel objects, suggesting that a greater population of hcrt neurons is being recruited in response to social novelty.

Earlier work has implicated that hcrt signaling plays a role in social behavior. Accordingly, male *Hcrt receptor 1* gene (*Hcrtr1^-/-^*) null mutant mice exhibit reduced sociability and decreased preference for social novelty compared to wild type mice (Abbas et al., 2015). Male orexin null mutant mice show diminished behavioral response to an intruder in a resident-intruder paradigm (Kayaba et al., 2003). A microdialysis study in human epileptic patients found maximal hcrt release during social interactions (Blouin et al., 2013). While orexin/ataxin-3-transgenic mice with hcrt cell loss show deficits in social memory (Yang et al., 2013). Supported by these studies, our data suggest a novel role for hcrt neuron activity states in the encoding of social discrimination. We also report a significant correlation between initial social approach and interaction-induced hcrt neuron activity and the subsequent time mice spent engaging with stranger conspecifics, suggesting that the initial approach-induced engagement of the hcrt neurons determines the duration of subsequent social interaction.

While both females and males show greater hcrt activity in response to social interaction, we obtained sex differences in the extent of hcrt activity and social interaction in females compared with males. Overall, both the hcrt Ca^2+^ signal and amount of time spent in social sniffing habituated faster in females throughout the reciprocal social interaction test. The faster habituation of the hcrt neuron activity during social interaction in females may reflect a disengagement of this system during the initial stages of interaction. A difference in sex-dependent neural and social engagement is not surprising given the variable strategies utilized by female and male mice in social behavior (Malloy et al., 2005; Okuyama et al., 2016; Rao et al., 2019). Yet, while the optogenetic inhibition of hcrt neurons resulted in a deficit in the normal behavior of male mice towards conspecifics, this did not lead to a change in sociability of females. Pharmacological blockade of hcrt-1 receptors similarly disrupted social interaction in males only, suggesting that sex-dependent hcrt neuron-induced modulation of social behavior is hcrt dependent.

The difference in the activity phenotype of hcrt neurons may represent sex-specific differences in local LH hcrt neuron network dynamics, which aligns with the sex differences observed in other hypothalamic cell subpopulations (K. Hashikawa et al., 2017). The mechanisms underlying the sex differences may also be in part be due to the differential stress response that may be induced by same-sex social interactions in females and males. As social interactions may be more rewarding to females than males (Borland et al., 2019), the extent of the engagement of hcrt neurons in same-sex social interaction may be dependent on the positive or negative valence to a social stimulus. In contrast, same-sex social interaction may be more stressful for males due to their increased tendency to show aggressive behaviors when interacting with other males (Malloy et al., 2005). Interestingly, the hcrt/orx system is highly regulated by stress in a sex-dependent manner (Grafe and Bhatnagar, 2020). Although mice were not exposed to stress prior to social interaction in our experiments, the capability of females for social buffering acute stress responses (Sterley et al., 2018; Taylor et al., 2000) may be a potential determinant in the differential engagement of neural populations based on the valence of the behavioral task. Interestingly, inhibition of the paraventricular CRH neurons (that are under the excitatory regulation of the hcrt system) (Bonnavion et al., 2015), similarly decreases social interaction in male mice (Sterley et al., 2018). Altogether, the sex-dependent recruitment of hcrt neurons in social behavior may have important implications for disorders associated with social deficits and that are more prevalent in men compared with women (Silber et al., 2002; Werling and Geschwind, 2013).

Hcrt neurons are key regulators of sleep, wakefulness and arousal (Carter et al., 2012; de Lecea, 2021; Inutsuka and Yamanaka, 2013; Sakurai, 2005; Tyree et al., 2018). Increased hcrt activity reduces sleep to wake transitions (Adamantidis et al., 2007; Carter et al., 2009) while reduced hcrt signaling or ablation of hcrt neurons has been associated with deficits in wakefulness and narcolepsy, respectively (Hara et al., 2001; Peyron et al., 2000; Sasaki et al., 2011; Tsunematsu et al., 2011). Decreased arousal and wakefulness may contribute to a reduced interest for social interaction. However, we did not find a change in locomotor activity upon acute hcrt neuron inhibition or pharmacological blockade of hcrt-1 receptors, suggesting that reduced interaction with conspecifics is not simply due to sedation. Given the strong association of narcolepsy and social impairments (Morse and Sanjeev, 2018; Quaedackers et al., 2019), a correlation between potential changes in sleep/wakefulness parameters and social interaction upon acute hcrt inhibition will provide more insight. Due to the involvement of hcrt in stress response (Grafe et al., 2018, 2017; Sargin, 2019), manipulation of hcrt neuron activity may interfere with generalized anxiety states. However, we did not observe changes in general anxiety-like behavior with acute hypocretin inhibition or hcrt-1 receptor antagonism, suggesting that the association between reduced hcrt activity and social interaction may not be attributed solely to changes in anxiety levels.

Hcrt neurons are located exclusively in the hypothalamus. However, they are a heterogenous group of cells with distinct neurochemical and electrophysiological characteristics (Mickelsen et al., 2017; Sagi et al., 2021). A substantial number of hcrt neurons co-express other neuropeptides and transmitters including dynorphin (Chou et al., 2001), and glutamate (Henny et al., 2010; Mickelsen et al., 2019; Schöne et al., 2012). Our experiments involving the blockade of hcrt-1 receptors revealed a disruption in social interaction, thereby strengthening the involvement of hcrt release in mediating social behavior. Further research is necessary to comprehensively understand how the additional peptides and transmitters co-released by hcrt neurons may potentially regulate social behavior. Recent research identified the expression of sex-specific genes, *Ddx3y* and *Xist* in distinct clusters of hcrt neurons, suggesting a role for sexual dimorphism in these neurons (Mickelsen et al., 2019). The specific role of these genes in hcrt-mediated sex differences and the extent to which these hcrt clusters contribute to social interaction remain to be determined. Furthermore, the bulk of our understanding of hcrt signaling has been derived from studies primarily focused on males. In our experiments, we did not consider the estrous cycle due to potential stress-related effects of the procedure as well as our experimental plan to include a randomized group of females. The comparison of variances using Levene’s test in key experiments did not show a significant difference between females and males, which indicates that sex-dependent differences in social interaction cannot be solely attributed to variations in estrous cycles. The absence of androgen and estrogen receptors in hcrt neurons adds another layer of complexity to the understanding of hcrt-mediated sex differences (Muschamp et al., 2007). Furthermore, it is noteworthy that ovariectomy has been found to have no discernible effect on the social interaction of females (Choi et al., 2022). This finding reinforces the importance of exploring sex-dependent mechanisms that underlie social behavior from alternative perspectives, allowing for a more comprehensive understanding.

To further investigate the mechanisms underlying the reduced hcrt signaling-induced disruption in social behavior of male mice, we performed cFos-dependent activity mapping in postsynaptic target regions that are targeted by hcrt neurons and are critical for social behavior. We identified that the activity of two key brain regions, the insular cortex and the VTA were significantly altered in male mice treated with the hcrt-1 receptor antagonist SB-334867 prior to social interaction. In contrast, cFos levels were comparable between vehicle-and SB-334867 administered female mice. These findings show that, unlike in males, reduced hcrt signaling in female mice does not result in alterations in social interaction or impact the social interaction induced activity of the examined brain regions. The insular cortex plays a pivotal role in governing emotional and interoceptive information (Gogolla, 2017; Wang et al., 2019), and is a vital component of the network responsible for social decision-making and social affect (Rogers-Carter et al., 2018). The VTA is also critical for motivated behavior, social reward and social approach (Gunaydin et al., 2014). Both the insular cortex and VTA receive strong hcrt inputs and express hcrt-1 receptors that modulate the activity of a variety of cell types in these regions (Hollander et al., 2008; Thomas et al., 2022). The differential sex-dependent changes in the activity of these regions point to potential circuits that are preferentially involved in hcrt-mediated social interaction. Both the insular cortex and VTA show disrupted connectivity in patients with autism spectrum disorders (ASD) and this is thought to be a contributing factor to the social impairments observed in these conditions (Caria and de Falco, 2015; Ebisch et al., 2011; Huang et al., 2021; Uddin and Menon, 2009). Given the higher prevalence of ASD in males, it is crucial to conduct further research on the sex-dependent modulatory role of hcrt connections to these regions during social behavior.

Overall, our findings provide a thorough characterization of hcrt neuron activity in social interaction and situate the hcrt system as a critical part of a larger network that plays an integral role in the modulation of social behavior. The observed sex differences further emphasize the need for the incorporation of sex as a biological variable in the design of studies focusing on the modulation of social behavior to understand the underlying mechanisms and circuits. Insights into the role of the hcrt system in social behavior will particularly be important in paving the way for novel treatment strategies targeting neuropsychiatric disorders associated with social deficits.

## Materials and Methods Animals

We used female and male mice (10 to 16-week old) *Hcrt^IRES-Cre^*mice on a C57BL6/J background. Age-, and sex-matched C57BL6/J mice were used as strangers. For vehicle or SB-334867 injections, C56Bl6/J mice were used. Mice were housed in standard housing conditions on 12 h light/dark cycle (light on at 07:00) with water and food available *ad libitum*. All experiments were conducted at the same time each day (12:00 – 16:00 pm). All experimental procedures were performed in accordance with the guidelines established by the Canadian Council on Animal Care and were approved by the Life and Environmental Sciences Animal Care Committee at the University of Calgary.

### Generation of the *Hcrt^IRES-Cre^* knock-in mouse line

*Hcrt^IRES-Cre^* mice were generated and gifted by Dr. Gina Leinninger. Briefly, a targeting vector for homologous recombination was generated by inserting an internal ribosome entry site (IRES)-Cre cassette into the 3’ end of the *hypocretin* mRNA product. A frt-flanked neo cassette was placed upstream of the IRES-Cre sequence. This *Hcrt^IRES-Cre^* targeting vector was linearized and electroporated into C57/Bl6 mouse embryonic stem cells. DNA from ES cell clones was analyzed via qPCR for loss of homozygosity using Taqman primer and probes for the genomic *Hcrt* insertion sites (*Hcrt:* Forward: TTTACCAAGAGACTGACAGCG, Reverse: CGGAGTCTGAACCCATCTTC, Probe: TCCTTGTCCTGATCCAAACTTCCCC). *NGF* was used as a copy number control (Soliman et al., 2007). Putative positive ES clones were expanded, confirmed for homologous recombination by Southern blot for neo and injected into blastocysts to generate chimeric mice. Male chimeras were bred with C57BL/6 albino females and produced germline progeny that express *Cre* and/or wild type *hcrt* (as assessed via PCR genotyping), confirming successful establishment of the *Hcrt^IRES-Cre^* line. (Forward Cre: CAC TGA TTT CGA CCA GGT TC, Reverse Cre: CAT CGC TCG ACC AGT TTA GT; Forward Hcrt wt: CTG GCT GTG TGG AGT GAA A, Reverse Hcrt wt: GGG GGA AGT TTG GAT CAG G. Cre band = 255 bp, Hcrt wt band = 460 bp).

*Hcrt^IRES-Cre^* mice were initially bred with *FlpO-Deleter* mice (Jackson Laboratories *B6.129S4-Gt(ROSA)26Sor^tm2(Flp*)Sor^/J*, stock #012930) to remove the frt-flanked neo cassette. Neo-deleted *Hcrt^IRES-Cre^*mice were then bred with C57/BL/6J mice (Jackson Laboratory, Strain #000664) to propagate the line.

### Stereotaxic Virus Injection and Optical Fiber Implantation

Mice were anaesthetized with 5% isoflurane before being transferred to a stereotaxic head frame (Kopf Instruments) and maintained at 2% isoflurane. Mice received analgesia (5 mg/kg meloxicam, s.c.) and fluid support (Lactated Ringer’s solution, s.c.) during surgery. Viral infusion was performed using a glass micropipette attached to a Nanoject III infusion system (Drummond Scientific) at 25 nl/sec for 10 sec. The infusion pipette was kept in place for 5 min after viral delivery and slowly elevated over 5 min. The following coordinates were used for virus injections: LH (from Bregma: AP -1.5 mm; ML ±1.0 mm; DV -5.2 mm). For fiber photometry experiments, mice received a unilateral infusion of AAV9-CAG-flex-GcAMP6s.WIRE.SV40 (Addgene viral prep # 100842-AAV9; 1.5 x 10^13^ vg/ml, 250 nl). For optogenetic experiments, mice received bilateral infusions of AAV2/9-EF1a-DIO-iC++-EYFP (Canadian Neurophotonics Platform Viral Core Facility RRID:SCR_016477; 1.7 x 10^13^ vg/ml, 250 nl per hemisphere) or AAV2/9-EF1a-DIO-EYFP (Canadian Neurophotonics Platform Viral Core Facility RRID:SCR_016477; 1.9 x 10^13^ vg/ml, 250 nl per hemisphere).

For fiber photometry experiments, viral infusions were followed by implantation of mono fiber optic cannulae (Neurophotometrics; 400 um, AP -1.5 mm; ML ±1.0 mm; DV -5.2 mm). For optogenetic experiments, dual fiber optic cannulae (Doric Lenses; 200 um, AP -1.5 mm; ML ±1.0 mm; DV -4.9 mm) were used. Implants were secured to the skull using super glue and dental cement. Behavior experiments were performed at least 3 weeks after surgery.

### Drug Injections

The hcrt receptor-1 antagonist SB-334867 (HelloBio) was dissolved in sterile water with 2% v/v DMSO and 10% w/v β-hydroxypropyl-cyclodextrin. SB-334867 or vehicle (2% v/v DMSO and 10% w/v β-hydroxypropyl-cyclodextrin in water) was injected intraperitoneally at a concentration of 30mg/kg 30 min prior to social interaction (Adamantidis et al., 2007; Ito et al., 2008; Thomas et al., 2022).

### *In vivo* Fiber Photometry Recording and Analysis

A Doric Lenses fiber photometry system with an LED driver and console was used to deliver the 405 nm isosbestic control and 465 nm excitation light during recordings. LEDs were connected to a Mini Cube filter set and a dichroic mirror array was used to permit simultaneous delivery of excitation light through a low autofluorescence mono fiber-optic patch cord to the fiber optic cannulae. The signal was detected by a photoreceiver (Newport Visible Femtowatt Photoreceiver Module Model 2151). A lock-in amplification mode controlled by the Doric Studio software was implemented (modulation frequencies: 208.6 Hz for 456 nm, 166.9 Hz for 405 nm, cut-off frequency for demodulated signal: 12 Hz). Light power was calibrated to deliver 30 µW light at the tip of the fiber using a photometer (Thor Labs).

The Doric Lenses fiber photometry console was set to trigger recording on remote delivery of a time-to-live (TTL) pulse from a separate computer running ANY-maze behavior capture software to time lock video and photometry recordings. Data from photometry and behavioral recordings were extracted and exported to MATLAB (MathWorks). Using custom MATLAB analyses, the 405 nm channel was used as an isosbestic control and the data recorded using this channel was fit to a biexponential decay. Data from the Ca^2+^-dependent 465 nm channel were then linearly scaled using the fitted decay of the isosbestic channel, correcting for any photobleaching which occurred during the recordings (Meng et al., 2018). The resulting vector was converted to a ΔF/F trace as previously described (Evans et al., 2022), and each trace was normalized as a z-score (Reichenbach et al., 2022). Z-scored ΔF/F traces were analyzed for area under the curve, peak frequency, and maximum fluorescence.

For social behavior recordings in home and novel cages, mice were first habituated to the fiber for 5 min. Fiber photometry analysis was conducted based on the baseline of 20 sec prior to the initiation of the social behavior test. 20 sec baseline period was selected to avoid the effects of the activity changes that can take place within a longer time window and may mask social interaction-induced changes that take place minutes after. Area under the curve was operationally defined as the summed area between the X-axis and the z-scored ΔF/F trace. Comparisons of AUC during specific behaviors were normalized by the summed duration of the behavioral epochs of interest. Peak frequency was refined using a peak detection filter of 2 standard deviations above the median z-scored ΔF/F value of the recording session (Evans et al., 2022; Moda-Sava et al., 2019). For all behavior-specific analyses, photometry and behavioral timeseries vectors were aligned using nearest-neighbours approximation to account for differences in framerates across recording modalities.

### *In vivo* Optogenetic Inhibition

Mice expressing EYFP (AAV2/9-EF1a-DIO-EYFP) or the inhibitory opsin iC++ (AAV2/9-EF1a-DIO-iC++-EYFP) were used for optogenetic inhibition experiments. Laser driver (Opto Engine LLC) was connected to a 1 x 2 fiber optic rotary joint (Doric Lenses) attached to a dual fiber-optic patch cord. Mice were habituated to the patch cord for 5 min prior to starting the behavior experiments. Blue light (470 nm, 17 mW) was delivered continuously via ANY-Maze software-controlled AMi-2 optogenetic interface (Stoelting Co.) connected to the laser driver.

### Behavioral Testing

The order of behavior tests was counterbalanced to control for any effect of testing order. Before each test, the area around and under cages or the surfaces of the apparatus were sprayed with 70% ethanol and wiped down with paper towel to reduce any residual odour. Prior to testing, mice were handled and habituated to the fiber optic patch cord for 4 days.

### 3-Chamber Social Interaction Test

The 3-chamber social interaction test was performed in two-stages in an acrylic 105 cm X 45 cm apparatus containing two plexiglass dividers, splitting the chamber into three equal areas. During the habituation stage, each mouse was first placed in the middle chamber for 5 min. The removal of the dividers allowed mice to freely explore the chamber for an additional 5 min. During the test stage, one area contained a sex-and age-matched C57BL/6J mouse confined in an inverted pencil cup (social zone) while the other area contained an empty cup (non-social zone). Mice were first placed in the middle chamber with dividers in place for 5 min. Dividers were then removed and mice were allowed to explore the chamber for 5 min. Time spent in each area was tracked and analyzed using ANY-Maze video tracking software (Stoelting Co.).

### Reciprocal Social Interaction Test

To examine reciprocal social behavior, resident experimental mice were kept in their home cage. Cage mates were separated for at least 15 min. For fiber photometry experiments, 5 min after being connected to the fiber optic patch cord, the cage mate (familiar mouse) of the experimental mouse was placed in the home cage and animals were allowed to freely interact for 5 min. 5 min after removal of the cage mate, a sex-and age-matched C57BL/6J mouse (stranger mouse) was placed in the home cage, and mice were allowed to freely interact for 5 min. For optogenetic experiments, the tests were spread across two consecutive days to reduce the likelihood of potential effects of prolonged light delivery and continual photoinhibition. The order of familiar or stranger mice introduced into the home cage of the experimental mice was counterbalanced across days. Blue light (470 nm, 17 mW) was turned on to allow 2 min of initial inhibition before the introduction of the familiar or stranger mouse. The experimental mouse and conspecific were allowed to freely interact for 5 min during continuous light delivery. Reciprocal social behavior in a novel cage was examined using the similar paradigm above in a novel mouse cage with new bedding material. Different stranger mice were introduced to mice tested in home cage and novel cage experiments. Multiple social encounters have been avoided to keep stranger mice unfamiliar. Different familiar mice were used in each day and cage mates were tested on different days. All videos were recorded using ANY-Maze video tracking software (Stoelting Co.).

### Social Behavior Analysis

We used the open-source automated positional tracking and behavioral classifications algorithms DeepLabCut (DLC; (Lauer et al., 2021) and Simple Behavior Analysis (SimBA; (Nilsson et al., 2020) to allow for unbiased, high-precision quantification and machine analysis of social behavior. A subset of social interaction trials was also manually scored by an experimenter blind to the groups to validate the resulting behavioral classifiers. Behavioral classifiers from SimBA quantified social behavior as the time the experimental mice engage in head-torso sniffing (snout of the experimental mouse was in direct proximity or direct contact with the head or torso of a conspecific) and anogenital sniffing (snout of the experimental mouse was in direct proximity to the anogenital region of a conspecific, including when actively following behind and sniffing a conspecific). Social novelty index was calculated as follows: (% time interacting with stranger – % time interacting with familiar) / (%time interacting with stranger + % time interacting with familiar).

#### Machine Scoring

For automated analysis of social behavior using behavioral classification algorithms, video recordings of social behavior experiments were first exported from ANY-maze to 20 frames-per-second MP4 video files. A subset of these video files was imported into DLC, where the experimenter blinded to the groups labelled the nose, left & right ears, left & right latissimus dorsi, tail base, and the centre of the torso for each animal present in the frames. These frames were then used by DLC to train the neural network to label the seven body parts described above. The trained neural network then analyzed all video recordings of social behavior experiments to track and plot the Cartesian coordinates of animals’ body parts for every frame of each video. This effectively provided the detailed spatial location of mice for every point in time throughout the experiment.

The results of the positional tracking analysis from DLC were exported to SimBA. We used a supervised-learning deep neural network within SimBA to create predictive classifiers for head-torso and anogenital sniffing. The resulting predictive classifiers were then used to automatically identify patterns in the positional data from DLC. Automated behavioral labels were validated for accuracy by reviewing visualizations of classification results produced by SimBA.

For verification, all results from SimBA were visualized and reviewed by the experimenter, and a subset of experiments were hand-scored to compare against the results of the automated analysis. The results of SimBA’s predictive classification for each experiment were exported for both analysis of behavior alone and combined analysis with the photometry recordings from the respective experiment. For photometry experiments, SimBA data and raw photometry traces were imported and aligned by a custom MATLAB script. For optogenetic experiments, SimBA data were analyzed and plotted.

#### Object Interaction Test

Mice were made familiar with an object (a small plastic puzzle piece) placed in their home cage for 72 h prior to the day of testing. A round glass marble was used as a novel object. 5 min after being connected to the fiber optic patch cord, novel or familiar object was introduced into the home cage of the experimental mouse. Each mouse was allowed to interact with the objects for 5 min. Videos were tracked and the time spent with each object was analyzed using ANY-Maze software (Stoelting Co.).

#### Open Field Test

Mice were placed in an acrylic 40 cm x 40 cm apparatus and allowed to explore for 5 min. Blue light (470 nm, 17 mW) was kept on continuously while the animal was in the open field. Videos were recorded and the total distance traveled, and the time spent in the outer and inner zones were analyzed using ANY-Maze video tracking software (Stoelting Co.).

#### Elevated Plus Maze

Mice were placed in the center of a plus-shaped apparatus consisting of two closed (36 cm) and two open (36 cm) arms. Blue light (470 nm, 17 mW) was kept on continuously while mice were allowed to explore for 5 min. Videos were recorded and the total distance traveled, and the time spent in each arm were analyzed using ANY-Maze video tracking software (Stoelting Co.).

#### Slice Electrophysiology

400 μm coronal slices comprising the LH were obtained in ice-cold oxygenated sucrose-substituted artificial cerebrospinal fluid (ACSF) using a Leica VT1000 S vibratome. Slices were recovered in aCSF (128 mM NaCl, 10 mM D-glucose, 26 mM NaHCO3, 2 mM CaCl2, 2 mM MgSO4, 3 mM KCl, 1.25 mM NaH2PO4, pH 7.4) for a minimum of 2 hours. During recovery and recordings, slices were saturated with 95% O2/5% CO2 at 30°C. Recording was performed in oxygenated aCSF flowing at a rate of 3-4 ml/min. The internal patch solution contained 120 mM potassium gluconate, 10 mM HEPES, 5 mM KCl, 2 mM MgCl2, 4 mM K2-ATP, 0.4 mM Na2-GTP, 10 mM Na2-phosphocreatine, with pH adjusted to 7.3. Neurons were visualized using IR-DIC on an Olympus BX51WI microscope and hcrt neurons were identified based on the expression of EYFP. Whole-cell recordings were obtained in current clamp mode using a MultiClamp 700B amplifier (Molecular Devices). Optogenetic inhibition through activation of iC++ was elicited using CoolLED pE-300^ultra^ illumination system at 470 nm. Data were filtered at 4 kHz and digitized at 20 kHz using Digidata 1550B and Clampex software (Molecular Devices).

#### Histology, Immunohistochemistry and Imaging

Mice were then deeply anaesthetized by isoflurane and transcardially perfused with 0.1 M PBS and 4% paraformaldehyde (PFA). Brains were extracted and post-fixed in 4 % PFA for 24 h and cryoprotected in 30% sucrose in 0.1 M PBS for 3-5 days until sunk. Brains were frozen and 40 µM thick coronal sections were obtained using a cryo-microtome (ThermoFisher) and collected in 6 series. Sections were stored in an antifreeze solution containing 30 % ethylene glycol, and 20 % glycerol in 0.1 M PBS and stored at −20°C for further analysis.

For immunolabeling, sections were washed three times in 0.1 M PBS for 10 min each. Sections were incubated with goat anti-hypocretin-1/orexin A antibody (1:500, Santa Cruz, sc-8070) and/or rabbit anti-melanin concentrating hormone (MCH) (1:500, Phoenix Pharmaceuticals), rabbit anti-GFP (1:500, Invitrogen) or with rabbit anti-cFos (1:1000, SYSY) antibodies diluted in 3 % normal donkey serum, 0.3 % Triton-X and 0.1 M PBS for 24 h. After three washes with 0.1 M PBS for 10 min each, sections were incubated with donkey anti-goat Alexa Fluor 594 (1:500, Jackson ImmunoResearch) and/or anti-goat Alexa Fluor 647 (1:500, Invitrogen), or anti-rabbit Alexa Fluor 647 (1:500, Invitrogen) diluted in 0.1 M PBS for 24 h. After three 10 min washes with 0.1 M PBS, sections were mounted on glass slides and coverslipped with PVA-DABCO mounting medium. Sections immunostained for cFos were were incubated with DAPI (1:500, HelloBio) in PBS for 15 min and washed three times in 0.1 M PBS before mounting. Sections were imaged using an Olympus FV3000 confocal microscope. Images were acquired with a 10X and 60X oil immersion objectives, with the aperture set to 1 airy unit. Z-stack image sequences were maximum intensity projected as a 2D image using Fiji. 2-3 sections comprising the LH were analyzed per mouse. Quantification of cells labeled with GFP and hypocretin-1 was performed using Fiji image processing software. Only mice with accurate stereotaxic targeting and viral expression were included in the analyses. For cFos analysis, sections were scanned at 10X magnification using a slide scanner (OLYMPUS VS120-L100-W; Richmond Hill, ON, CA). cFos positive cells were segmented from background based on fluorescent intensity and label morphology using *Ilastik* (Berg et al., 2019). Images were then registered to the Allen Brain Institute mouse brain reference atlas to assess the expression density of segmented cFos positive cells using FASTMAP (Terstege et al., 2022). Cells in the following regions were quantified: Insular cortex, infralimbic cortex, prelimbic cortex, CA1 region of the hippocampus, lateral amygdala, bed nucleus of the stria terminalis (BNST), nucleus accumbens (NAc), and the ventral tegmental area (VTA). The expression density was calculated as the number of cFos positive cells divided by the area of the region for each brain section.

### Statistical Analysis

Statistical analysis was performed using GraphPad Prism 9, except three-way ANOVAs which were conducted using Statistica software. Tukey’s multiple comparisons tests were applied when the F test for the interaction is significant (Rosner, 2000). In the absence of a significant interaction, we planned a priori comparisons between the groups and separately analyzed these comparisons using Fisher’s LSD test. These differences are indicated on the graphs with an octothorpe signal. In cFos mapping experiments, Bonferroni adjustments were used to reduce the risk of Type 1 error with the large number of regions analyzed. The AUC of the Ca2+ signal from one mouse in Figure 1 was identified to be an outlier (Grubb’s test; alpha = 0.05, G= 1.77, outlier Y = 533.7) and removed from the analysis. All statistical data are reported in Supplementary Tables S1-S5.

## Acknowledgements

We thank the transgenic cores at University of Michigan and Van Andel Research Institute for assistance generating *Hcrt^IRES-Cre^* mice. David P. Olson graciously shared the Cre-inducible Rosa^eGFP-L10a^ reporter line used in these studies. Funding for this study was provided by an NSERC Discovery Grant (RGPIN-2020-05305) and an Arrell Family Foundation Future Leader in Canadian Brain Research grant to DS, an NSERC Discovery Grant (RGPIN-2018-05135) to JRE. DJT received fellowships from NSERC and the Canadian Open Neuroscience Platform. DP received Harley Hotchkiss-Samuel Weiss Postdoctoral Fellowship.

## Author contributions

D.S. and M.D. designed the experiments and wrote the manuscript. R.B. and G.M.L. designed and made the mouse line. N.J. and M.T. performed the surgeries. M.D. and D.P. performed the behavioral experiments. M.D. performed the histological procedures. D.S., M.D., D.J.T. and J.R.E. conducted the analysis.

## Data availability

All data that support the findings of this study are present in the manuscript and the supplementary materials, and are available from the corresponding author upon request.

## Code availability

All custom scripts used for photometry analysis are publicly available in a GitHub repository (https://github.com/dterstege/PublicationRepo/tree/main/Dawson2022).

