## Supplemental Figures for "Hypocretin/orexin neurons encode social discrimination and exhibit a sex-dependent necessity for social interaction"

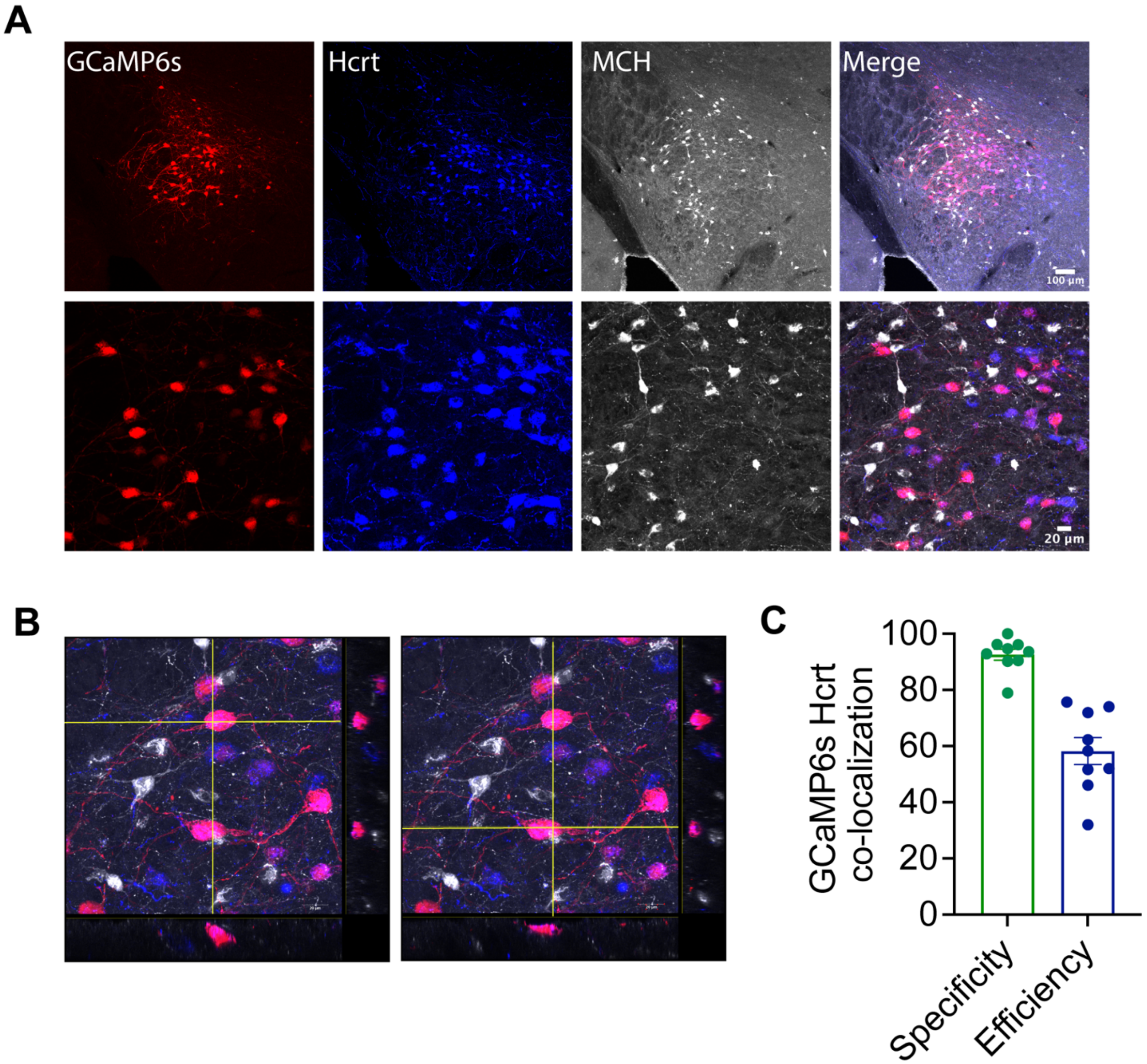


**Figure S1.** ***Hcrt*$^{IRES\text{-}Cre}$ line validation. A)** Representative images showing co-localization of AAV-CAG-flex-GCaMP6s expression (red) and Hcrt immunostaining (blue) or melanin-concentrating hormone (MCH) immunostaining (gray) in the LH. Zoomed images are shown below. **B)** Cells co-expressing GCaMP6s and Hcrt are negative for another large LH cell population, MCH immunoreactivity. Below: xz plane; right: yz plane are shown. **C)** Quantification of specificity (% virus-labeled cells expressing Hcrt) and efficiency (% Hcrt cells co-expressing the virus) from n=9 mice (4 males, 5 females).

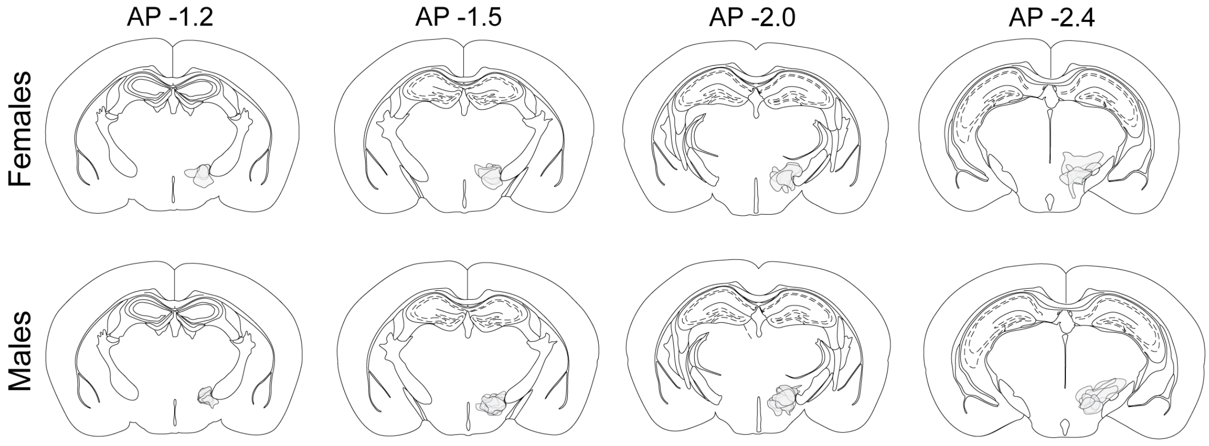


**Figure S2. Spread and validation of GCaMP6s expression in female and male mice.** GCaMP6s expression was restricted to AP -1.2 to 2.4. Gray areas denote the viral spread from 7 female and 7 male in *Hcrt*$^{IRES\text{-}Cre}$ mice infused with AAV9-CAG-flex-GcAMP6s.WIRE.SV40.

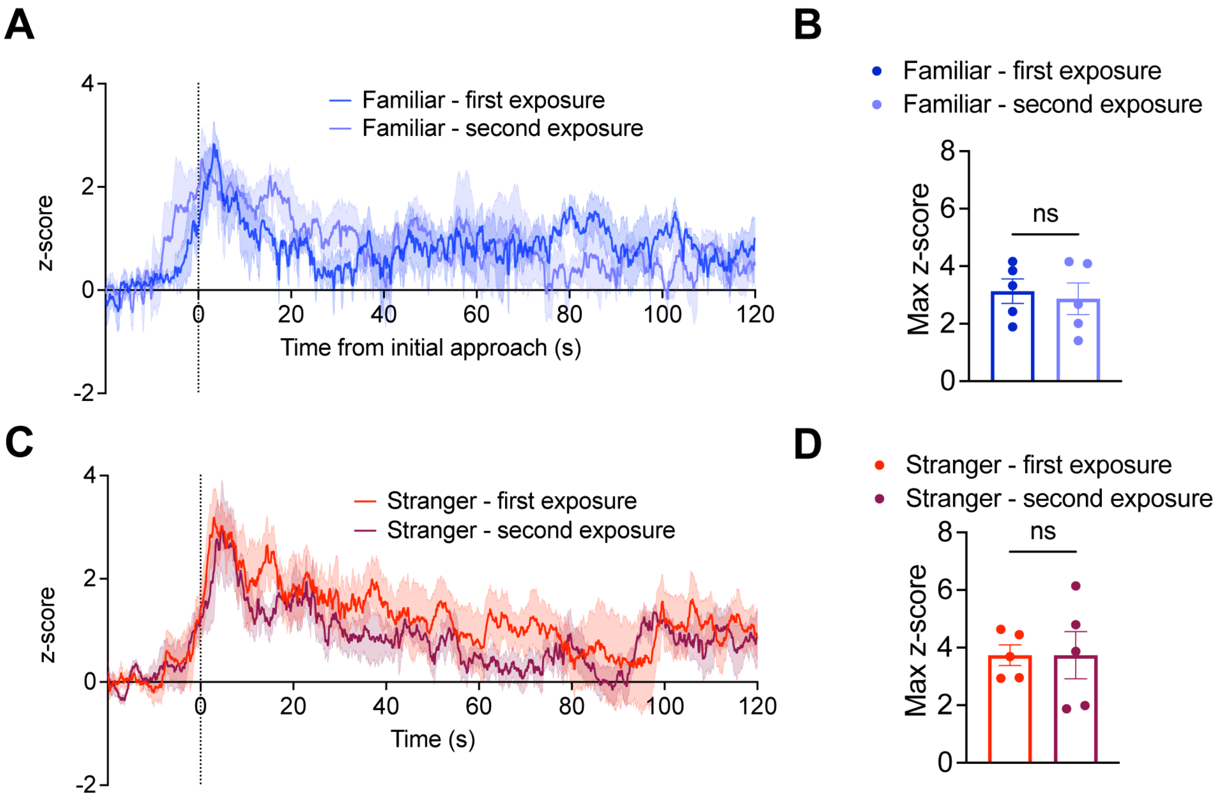


**Figure S3. Hcrt neuron activity remains the same after subsequent interactions with familiar and stranger conspecifics. A)** Averaged $Ca^{2+}$ signal changes in hcrt neurons in mice during subsequent exposures to a familiar mouse. After an initial 5 min interaction and 5 min baseline recording alone in the cage, the familiar conspecific was re-introduced into the home cage for 5 more min. **B)** The maximum hcrt $Ca^{2+}$ signal in response to initial approach and sniffing is comparable between subsequent exposures to a familiar cage mate (paired t-test: $p = 0.51$). **C)** Averaged $Ca^{2+}$ signal changes in hcrt neurons in mice during subsequent exposures to a stranger mouse. After an initial 5 min interaction and 5 min baseline recording alone in the cage, the stranger conspecific was re-introduced into the home cage for 5 more min. **D)** The maximum hcrt $Ca^{2+}$ signal in response to initial approach and sniffing is comparable between subsequent exposures to a stranger conspecific (paired t-test: $p = 0.99$). Data represent mean ± SEM.

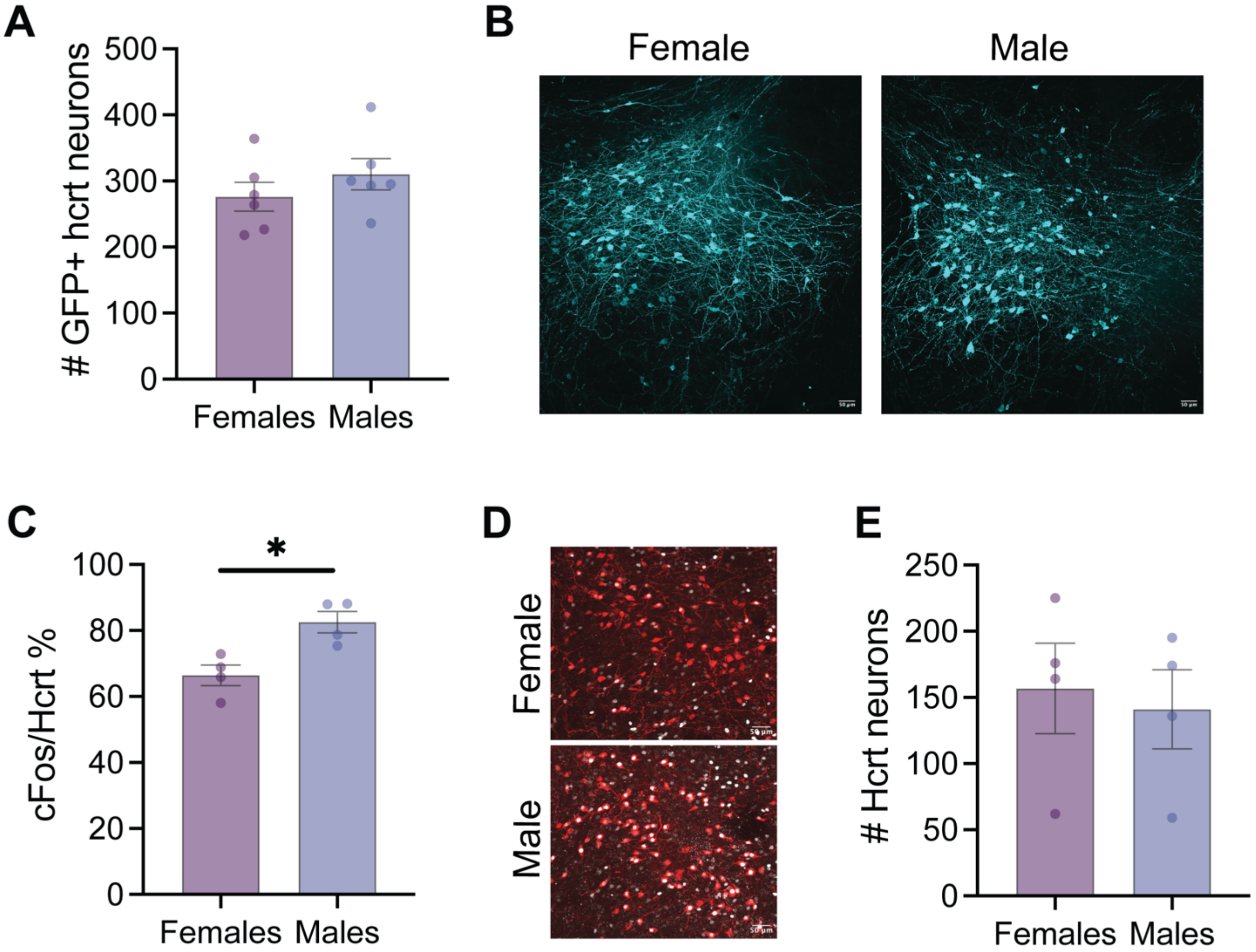


**Figure S4. Larger percentage of Hcrt neurons express cFos after social interaction in male mice, compared with females without a difference in the number of virus-infected cells or hcrt neurons. A)** The number of LH hcrt neurons infected with AAV encoding GCaMP6s is similar between female and male mice (unpaired t-test, t = 1.05, df = 10, $p$ = 0.32). **B)** Representative images showing GFP expression in the LH of Hcrt-cre female and male mice infected with AAV9-CAG-flex-GcAMP6s.WIRE.SV40. **C)** Percentage of hcrt neurons expressing cFos after social interaction is significantly greater in male mice, compared with females (unpaired t-test, t = 3.56, df = 6, *$p$ = 0.012). **D)** Representative images showing hcrt (red) and cFos (gray) immunostaining in the LH. **E)** The number of hcrt positive neurons in the LH is similar between female and male mice (males (unpaired t-test, t = 0.34, df = 6, $p$ = 0.74). Data represent mean ± SEM.

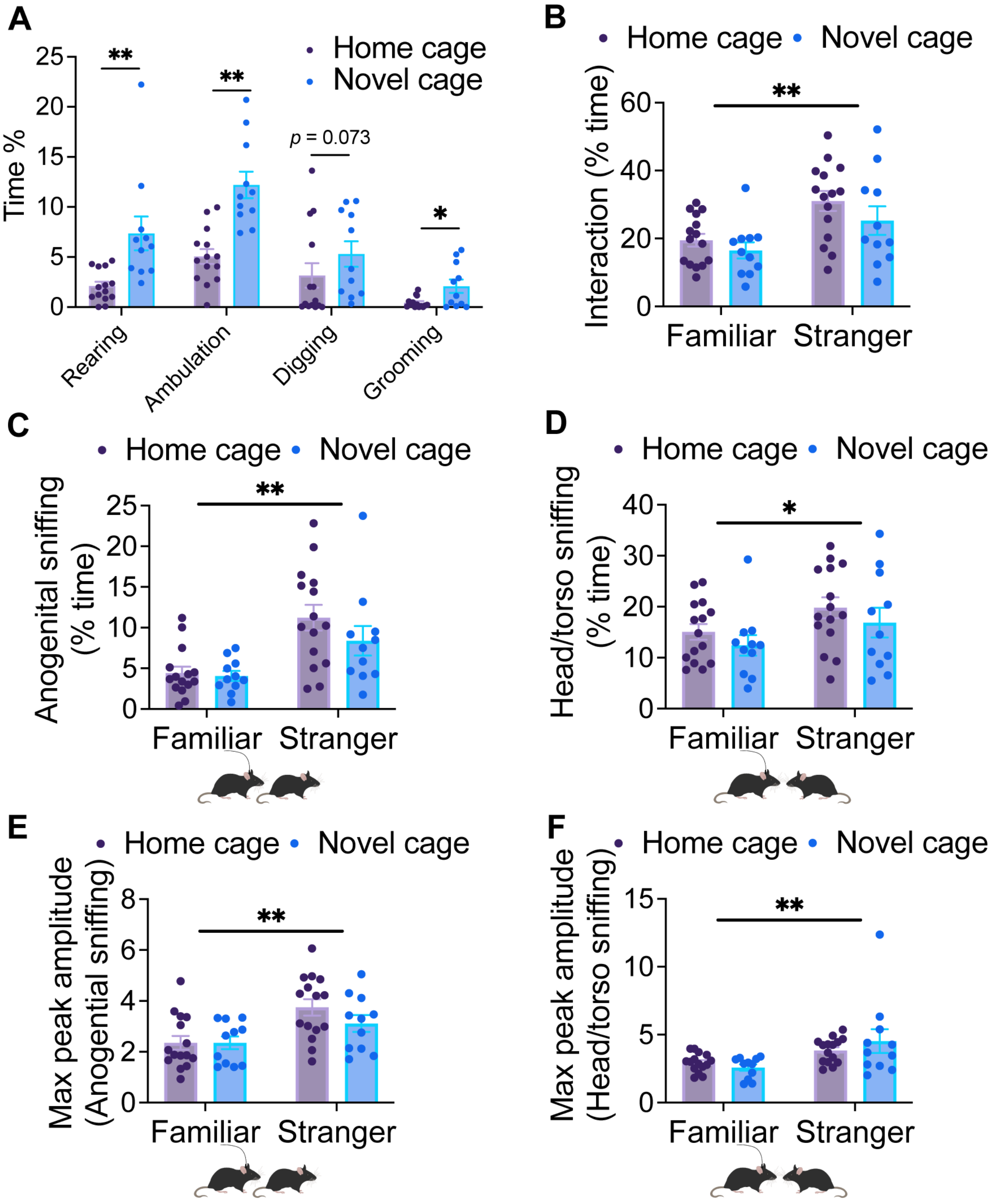


**Figure S5. Comparison of social sniffing behavior and hcrt photometry signal between home cage and novel cage. A)** Mice spent greater time engaging in non-social behaviors including rearing, ambulation and grooming in the novel cage (n=11 mice) compared to the home cage (n=14 mice) (unpaired t-test; $*p < 0.05$, $**p < 0.01$). **B)** Comparison of time mice spent interacting with familiar and stranger conspecifics in home and novel cages (two-way ANOVA; effect of cage: $F(1, 48) = 2.28$, $p = 0.14$, effect of conspecific: $F(1, 48) = 12.33$, $**p = 0.001$). **C)** Comparison of time mice were engaged in anogenital sniffing of familiar and stranger conspecifics in home and novel cages (two-way ANOVA; effect of cage: $F(1, 48) = 1.48$, $p = 0.23$, effect of conspecific: $F(1, 48) = 17.91$, $**p = 0.0001$). **D)** Comparison of time mice were engaged in head/torso sniffing of familiar and stranger conspecifics in home and novel cages (two-way

ANOVA; effect of cage: $F(1, 48) = 1.69$, $p = 0.2$, effect of conspecific: $F(1, 48) = 4.65$, *$p = 0.036$). **E)** Comparison of the maximum activity peaks of hcrt neurons during anogenital sniffing in home and novel cages (two-way ANOVA; effect of cage: $F(1, 48) = 1.1$, $p = 0.3$, effect of conspecific: $F(1, 48) = 12.53$, **$p = 0.0009$). **F)** Comparison of the maximum activity peaks of hcrt neurons during head/torso sniffing in home and novel cages (two-way ANOVA; effect of cage: $F(1, 48) = 0.15$, $p = 0.7$, effect of conspecific: $F(1, 48) = 11.42$, **$p = 0.0015$). Data represent mean ± SEM.

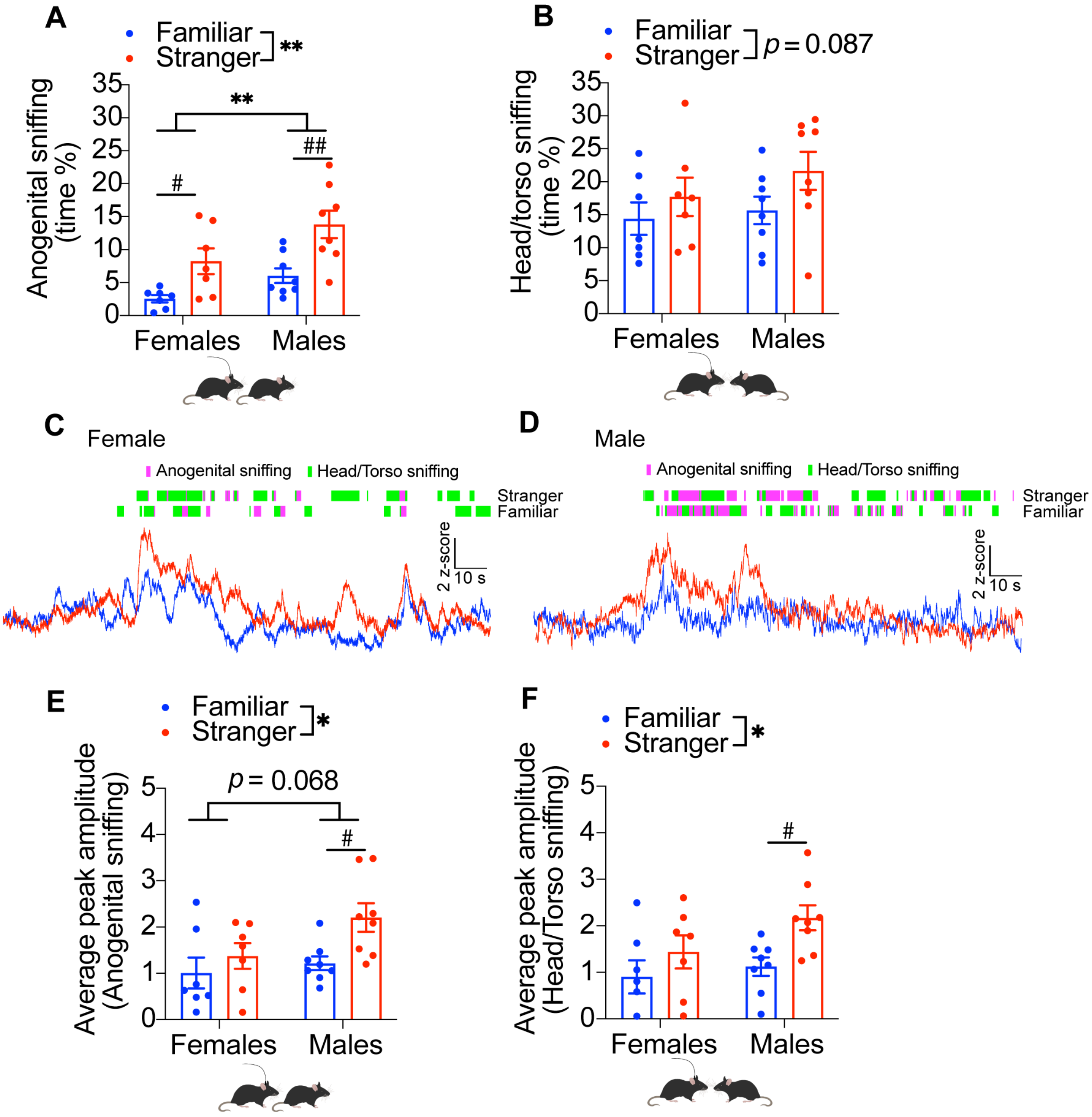


**Figure S6. Social behavior and hcrt neuron activity during anogenital and head/torso sniffing in female and male mice. A)** Female and male mice spent greater time engaged in anogenital sniffing during interaction with stranger conspecifics compared to familiar (two-way ANOVA; effect of conspecific: $F(1, 26) = 18$, $^{**}p = 0.002$, effect of sex: $F(1, 26) = 8.19$, $^{**}p = 0.0082$, Fisher's LSD; familiar vs stranger: females, $^{\#}p = 0.02$, males, $^{\#\#}p = 0.001$). **B)** The time female and male mice spent engaged in head/torso sniffing during interaction with stranger conspecifics compared to familiar was larger but did not reach significance (two-way ANOVA; effect of conspecific: $F(1, 26) = 3.17$, $p = 0.087$, effect of sex: $F(1, 26) = 0.99$, $p = 0.33$). Representative photometry traces of hcrt neurons from a female **(C)** and a male **(D)** mouse are shown. Interaction bouts corresponding to anogenital and head/torso sniffing time locked to the photometry are shown above. **E)** The average amplitude of hcrt neuron activity peaks during anogenital sniffing (average of 10 sniffing bouts) in female and male mice is greater during interaction with stranger conspecifics compared with familiar (two-way ANOVA; effect of conspecific: $F(1, 26) = 6.16$, *$p = 0.02$, effect of sex: $F(1, 26) = 3.62$, $p = 0.068$, Fisher's LSD; familiar vs stranger: females, $p = 0.36$, males, $^{\#}p = 0.013$). **F)** The average amplitude of hcrt neuron activity peaks (average of 10 sniffing bouts) during head/torso sniffing in female and male mice is greater during interaction with stranger conspecifics compared with familiar (two-way ANOVA; effect of conspecific: $F(1, 26) = 7.21$, *$p = 0.012$, effect of sex: $F(1, 26) = 2.6$, $p = 0.12$, Fisher's LSD; familiar vs stranger: females, $p = 0.22$, males, $^{\#}p = 0.015$). Data represent mean ± SEM.

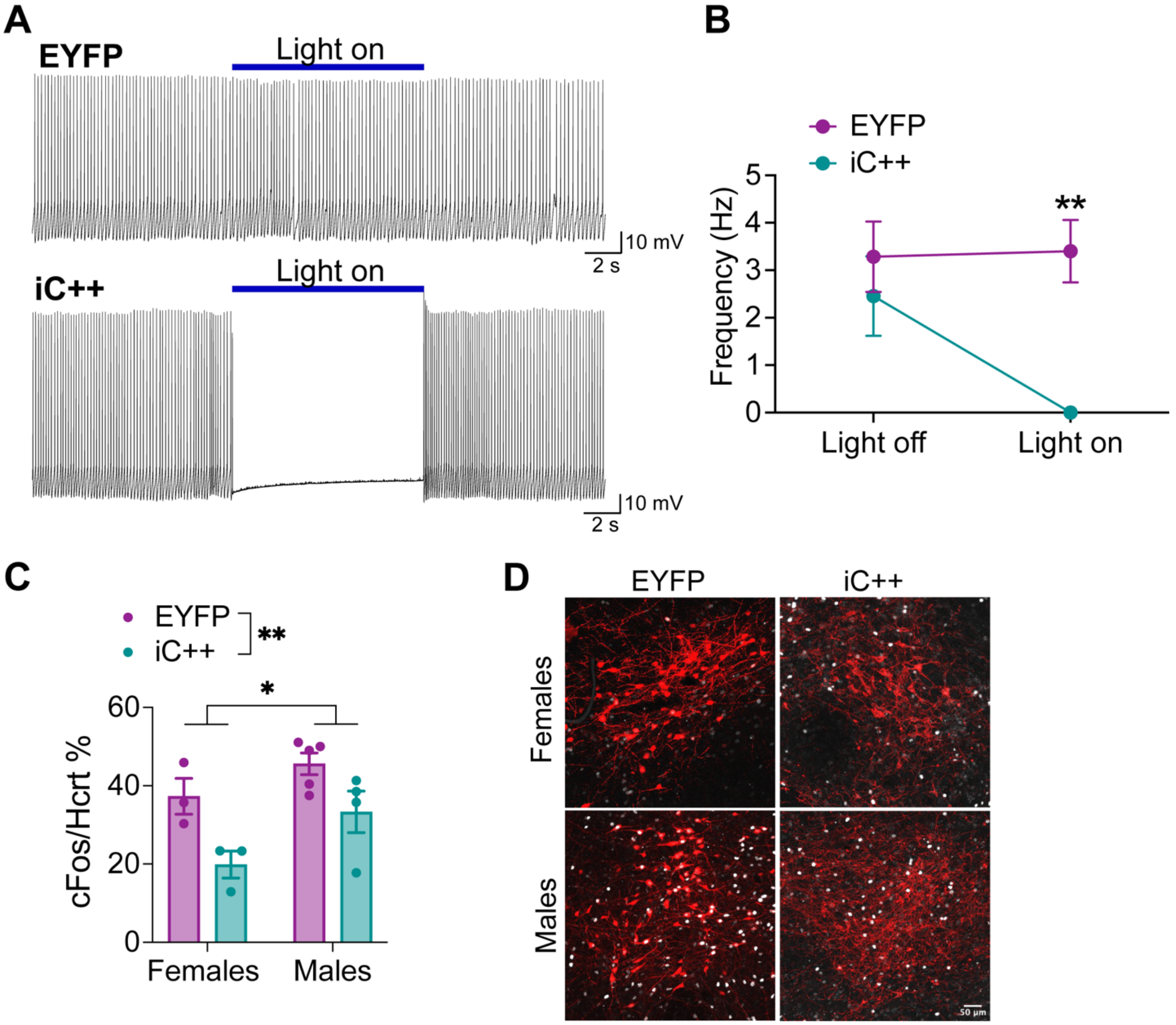


**Figure S7. Electrophysiological validation of photoinhibition of hcrt neuron activity. A)** Representative current clamp recordings of hcrt neurons from a EYFP- and iC++- expressing mouse. **B)** The effect of continuous blue light stimulation on the frequency of action potentials in hcrt neurons of EYFP and iC++ mice. 20-50 pA current was applied to keep neurons firing at 2-4 Hz (EYFP: n = 7 neurons, two female mice; iC++: n = 7 neurons, two female mice; two-way ANOVA: light x group interaction: $F(1, 12) = 8.56$, $*p = 0.013$, Tukey's multiple comparisons test; light off EYFP vs iC++ $p = 0.80$, light on EYFP vs iC++ $**p = 0.008$). **C)** The percentage of hcrt neurons expressing cFos after social interaction is decreased in mice expressing iC++, compared with those expressing EYFP (two-way ANOVA; effect of group: $F(1, 11) = 12.62$, $**p = 0.004$). cFos expression in hcrt neurons is greater in male mice, compared with females (two-way ANOVA; effect of sex: $F(1, 11) = 6.78$, $*p = 0.02$). Data represent mean ± SEM.

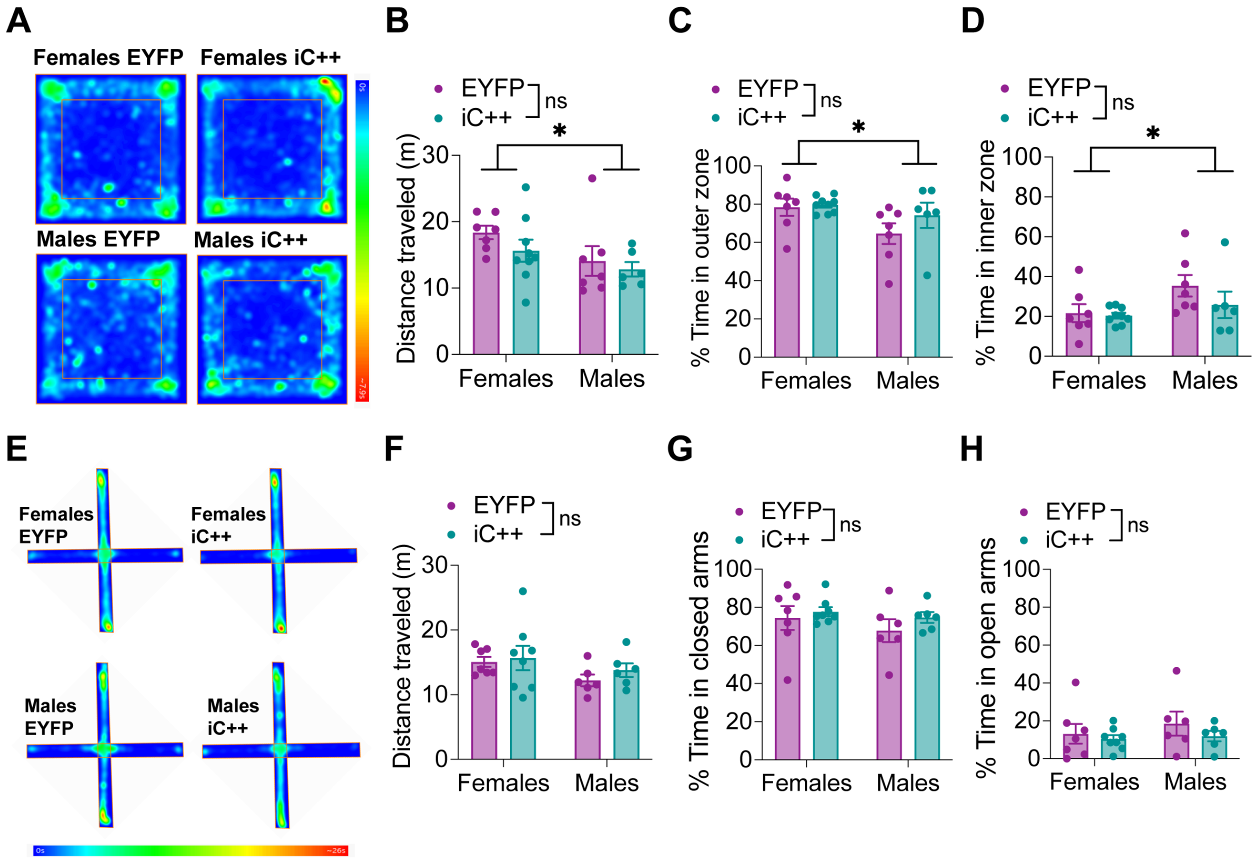


**Figure S8. Effect of acute optogenetic inhibition of hcrt neurons on locomotor activity and anxiety-like behavior. A)** Averaged heat maps showing the distance traveled in an open field in the presence of blue light in female and male EYFP and iC++ mice. B) Acute inhibition of hcrt neuron activity during open field test did not lead to changes in distance traveled. Females traveled greater distance than males (two-way ANOVA; effect of group: F(1, 25) = 1.46, *p* = 0.24, effect of sex: F(1, 25) = 4.62, **p* = 0.042). C) Acute inhibition of hcrt neuron activity during open field test did not affect time in the outer zone of the open field. Female mice spent greater time in the outer zone (two-way ANOVA; effect of group: F(1, 25) = 1.45, *p* = 0.24, effect of sex: F(1, 25) = 4.64, **p* = 0.041). **D)** Acute inhibition of hcrt neuron activity during open field test did not affect time in the inner zone of the open field. Female mice spent less time in the inner zone (two-way ANOVA; effect of group F(1, 25) = 1.45, *p* = 0.24, effect of sex: F(1, 25) = 4.64, **p* = 0.041). E) Averaged heat maps showing the distance traveled in an elevated plus maze in the presence of blue light in female and male EYFP and iC++ mice. Acute inhibition of hcrt neuron activity during elevated plus maze did not affect F) total distance traveled (two-way ANOVA; effect of group: F(1, 23) = 0.64, *p* = 0.43, effect of sex: F(1, 23) = 3.02, *p* = 0.09), G) time spent in closed arms (two-way ANOVA; effect of group: F(1, 23) = 1.21, *p* = 0.28, effect of sex: F(1, 23) = 1.07, *p* = 0.31) or H) time in open arms (two-way ANOVA; effect of group: F(1, 23) = 1.19, *p* = 0.28, effect of sex: F(1, 23) = 0.67, *p* = 0.42) of the elevated plus maze. Data represent mean ± SEM. ns, non-significant.

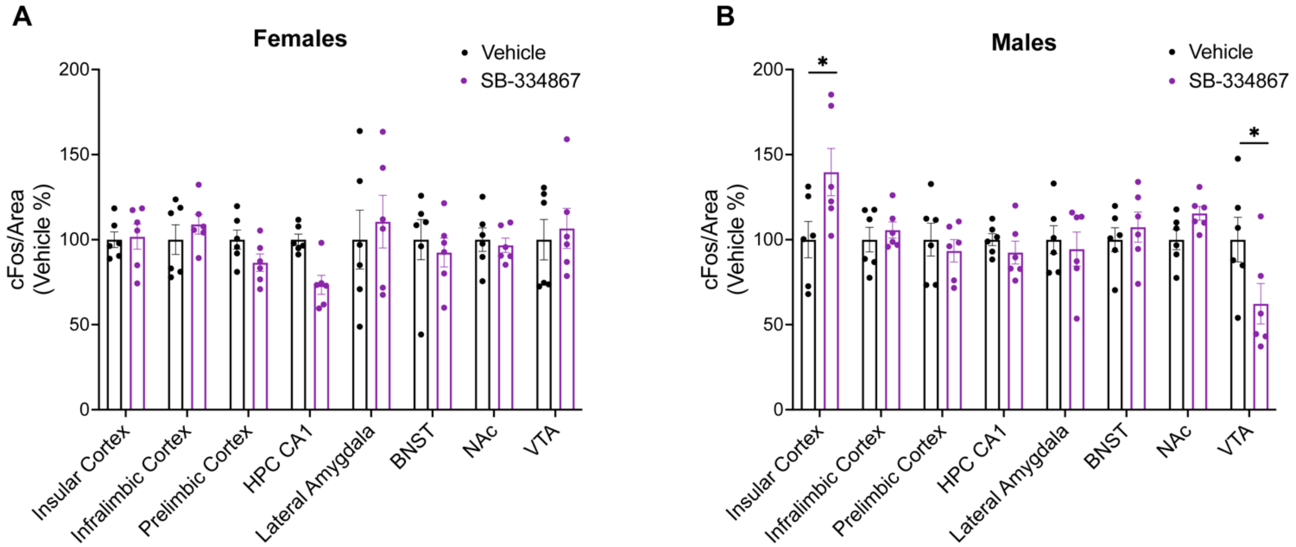


**Figure S9. Blocking hcrt-1 receptors during social interaction leads to increased insular cortex and decreased VTA activity in male mice.** Percent change in cFos density in mice injected with SB-334867 during social interaction relative to mice injected with vehicle. SB-334867-injected female mice show no changes **(A)** while SB-334867-injected male mice show significantly increased cFos density in insular cortex and reduced cFos density in the VTA, compared with vehicle-injected male mice **(B)**. HPC, hippocampus; BNST, bed nucleus of the stria terminalis; NAc, nucleus accumbens; VTA, ventral tegmental area.
